# PenTag, a Versatile Platform for Synthesizing Protein-Polymer Biohybrid Materials

**DOI:** 10.1101/2023.05.18.541264

**Authors:** Hasti Mohsenin, Jennifer Pacheco, Svenja Kemmer, Hanna J. Wagner, Nico Höfflin, Toquinha Bergmann, Tim Baumann, Carolina Jerez-Longres, Alexander Ripp, Nikolaus Jork, Henning J. Jessen, Martin Fussenegger, Maja Köhn, Jens Timmer, Wilfried Weber

**Affiliations:** Signalling Research Centres BIOSS and CIBSS, University of Freiburg, Freiburg, Germany; Faculty of Biology, University of Freiburg, Freiburg, Germany; INM ̶ Leibniz Institute for New Materials, Saarbrücken, Germany; Saarland University, Department of Materials Science and Engineering, Saarbrücken, Germany; Institute of Physics, University of Freiburg, Freiburg, Germany; Freiburg Center for Data Analysis and Modelling (FDM), University of Freiburg, Freiburg, Germany; SGBM - Spemann Graduate School of Biology and Medicine, University of Freiburg, Freiburg, Germany; Institute of Organic Chemistry, University of Freiburg, Freiburg, Germany; Cluster of Excellence LivMats, University of Freiburg, Freiburg, Germany; Department of Biosystems Science and Engineering, ETH Zurich and Faculty of Science, University of Basel, CH-4058 Basel, Switzerland

**Keywords:** covalent protein-ligand conjugation, covalent crosslinking, stimuli-responsive hydrogels, information-processing materials

## Abstract

The site-specific and covalent conjugation of proteins on solid supports and in hydrogels is the basis for the synthesis of biohybrid materials offering broad applications. Current methods for conjugating proteins to desired targets are often challenging due to unspecific binding, unstable (non-covalent) coupling, or expensive and difficult-to-synthesize ligand molecules. Here, we present PenTag, an approach for the biorthogonal, highly-specific and covalent conjugation of a protein to its ligand for various applications in materials sciences. We engineered penicillin-binding protein 3 (PBP3) and showed that this protein can be used for the stable and spontaneous conjugation of proteins to dyes, polymers, or solid supports. We applied PenTag as a crosslinking tool for synthesizing stimuli-responsive hydrogels or for the development of a biohybrid material system performing computational operations emulating a 4:2 encoder. Based on this broad applicability and the use of a small, cheap and easy-to-functionalize ligand and a stable, soluble recombinant protein, we see PenTag as a versatile approach towards biohybrid material synthesis.

## 1. Introduction

Response to environmental stimuli is a fundamental feature of living systems.[1] Implementation of this feature into materials systems results in the development of smart, stimuli-responsive materials that have applications in drug delivery, diagnostics, and biosensing.[2–4] Proteins, as powerful biological building blocks, equip materials with sensory, regulatory, and communicational functions.[5–8] By covalent coupling of proteins to their ligands, a platform for stable and robust anchoring of proteins to small molecules, solid supports and materials can be created.[9–13] Covalent attachment of chemicals and oligonucleotides to antibodies have resulted in antibody-drug conjugates (ADCs) and antibody-oligonucleotide conjugates (AOCs) and plays an important role as novel therapeutic vehicles for the treatment of various cancers and for non-oncological diseases.[14,15]

Several techniques have been developed for site-specific conjugation and modification of proteins through covalent binding that have paved their way through various applications in visualizing and manipulating biological processes.[16] For example for site-specific protein conjugation, cysteine residues in proteins can be used for bioconjugation of proteins via Thiol-Michael addition reactions of maleimides or vinylsulfones.[17–19] However, if more site-specificity is required, enzymatic labelling techniques, based on, for example *E. coli* biotin ligase BirA, or *Staphylococcus aureus* sortase A (SrtA)-mediated transpeptidation reactions are superior.[20,21] For instance, SrtA has been used for the development of protein-based materials for various applications.[9,22,23] These systems, however powerful, require addition of an additional enzyme that catalyses the coupling reaction, which need to be followed by the removal of the added enzymes or the by-products.[16] There are some protein tags that do not require an additional enzyme, for instance SNAP-, Halo-, and CLIP-Tags are three examples of self-labelling protein tags that can be used for labelling of proteins for diverse applications such as the specific immobilization and purification of proteins.[24–28] Such tags facilitate the downstream processing, as no further purification would be required. These tags are based on engineered self-modifying enzymes that spontaneously react with O^6^-benzylguanine (BG), chloroalkane, and benzylcytosine (BC) derivatives, respectively, leading to a covalent conjugation of the protein to its ligand. These tags are widely used for the spontaneous coupling of proteins to different ligands. For example, SNAP-Tag-mediated crosslinking have been used for the development of light-controllable hydrogels.[29] However, the required reactive groups are fine chemicals and available in small quantities at rather high prices. Amine-functionalized BG, BC, and HaloTag ligands cost 170, 600, and 140 USD per gram, respectively.[30]

In addition to mediating site-specific conjugation, an ideal protein tag for protein-based systems and biohybrid materials enables the detection, visualization, and purification of its fusion partner, confers increased solubility and is able to link the protein of interest to various types of molecules for example polymers and hydrogels. For some applications, it would be beneficial if the protein tag is small in size and forms a fast and stable bond at a variety of different temperatures and buffer conditions.[31,32] Additionally, an ideal ligand would be a simple and cheap chemical that is widely commercially available. Therefore, a novel conjugation technique is demanded that promises the following points: a simple, cheap, and easy-to-functionalize ligand, a recombinant soluble and solubilizing protein, a one-step spontaneous reaction that does not require the addition of further chemicals or biological molecules, nor needs the removal of by-product species, and finally a system that can be used orthogonally to the already existing systems without interfering with them.

ß-lactam antibiotics such as ampicillin have the interesting property that they bind specifically and covalently to penicillin-binding proteins (PBPs). PBPs are transpeptidase enzymes involved in bacterial cell-wall biosynthesis and are the target of ß-lactam antibiotics[33]. Interaction of PBPs with ß-lactam antibiotics starts with the formation of a non-covalent Henri-Michaelis complex and is followed by the attack of the serine residue in the active site of the enzyme to the ß-lactam ring, that results in a covalent acyl-PBP bond.[34,35] In previous works we have applied this specific coupling for the development of microfluidic, point-of-care- suitable assays for electrochemical antibiotics detection in various settings.[36–39]

Here, we exploit the spontaneous, covalent interaction between ampicillin and PenTag for the development of a protein tag and demonstrate its application in protein immobilization and purification. We use this conjugation technique as a building block towards the design of biohybrid materials performing fundamental computational operations, while showing its biorthogonality to the already existing systems. Besides broad application potential and ease of handling, the bulk availability of ampicillin at very low cost (1.6 USD per gram[30]) is an interesting factor in synthesizing larger quantities of biomaterials benefiting from this conjugation technique.

## 2. Results and Discussion

### 2.1. Engineered truncated PBP3 binds stably to ß-lactam antibiotics

Large proteins devour more metabolic energy during overexpression in *E. coli* compared to smaller proteins.[31] Further, large proteins represent a less favourable cost-benefit ratio if only one part of the protein structure is needed. However, reducing the size of proteins must be carefully planned and tested to avoid a negative impact on their functionalities. We therefore aimed at the design of a truncated version of PBP3, which is still able to bind to ß-lactam antibiotics. To this aim, we exploited a mutated PBP3[36,40] and the known crystal structure of PBP3[41] to truncate the C-terminal hydrophobic membrane anchor thereby generating a new version of PBP3, termed PBP3-t (**Figure** 1a). We produced hexahistidine-functionalized PBP3- t in *E. coli*, purified it by immobilized metal affinity chromatography (IMAC), and performed analysis by SDS-PAGE and Coomassie staining. The protein migrated at a position corresponding to 30.2 kDa, which is in agreement with its predicted molecular weight (**Figure** S1a).

**Figure 1.**
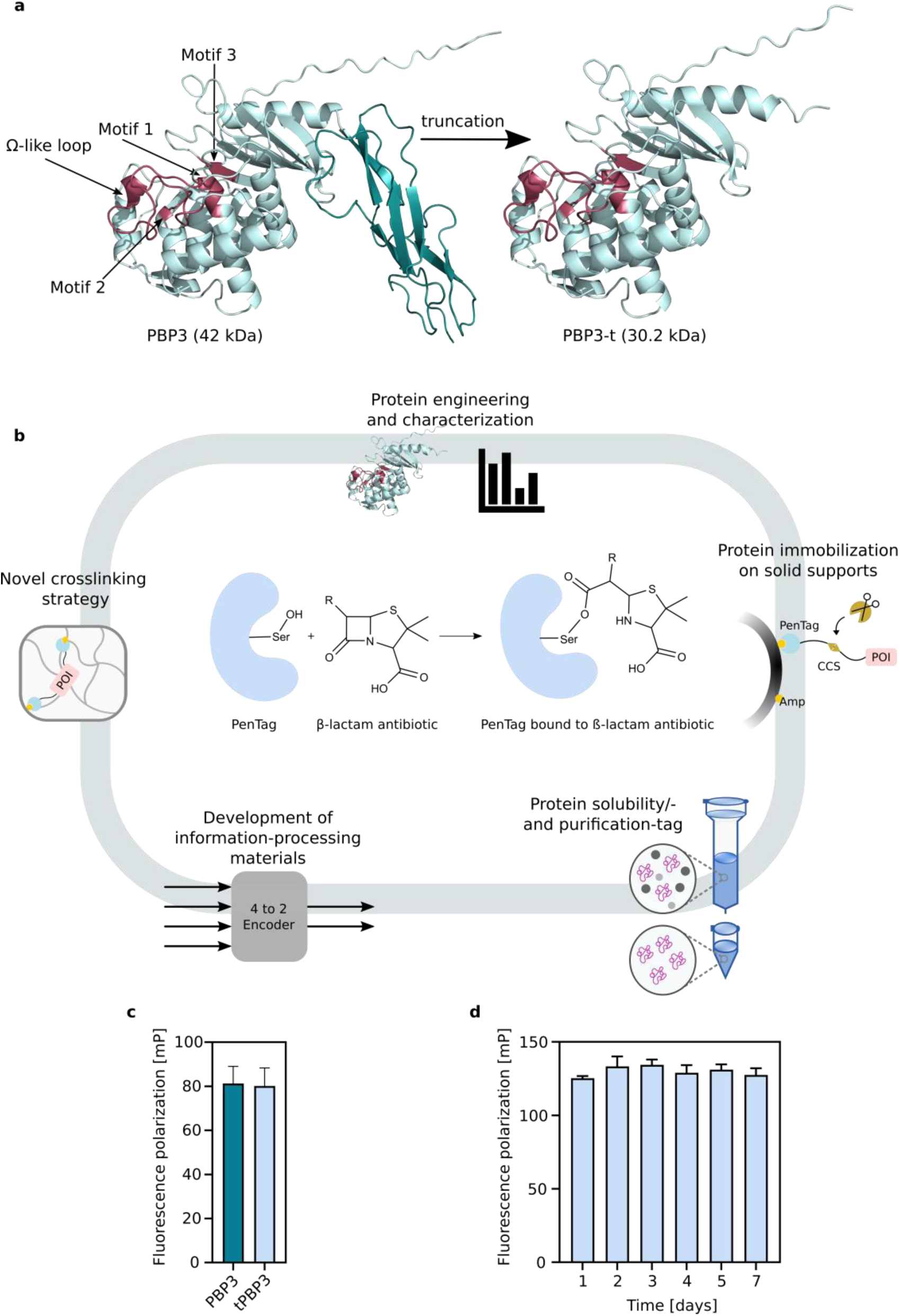
Engineering a new truncated version of PBP3. **a**, Structure of full-length PBP3 (with membrane anchor, left) and upon truncation (right). The membrane anchor is shown in dark cyan and the rest of the protein shown in light cyan. In red, the active site of the protein consisting of motif 1, 2, 3 and Ω-like loop is shown. Protein structure prediction is done by AlphaFold 2[44,45] and figures are generated by PyMOL[46]. **b**, Mechanism of conjugation of ß- lactam antibiotics to PenTag and an overview of applications that could benefit from this conjugation technique. **c,** Fluorescence polarization measurements to compare the PBP3/bocillin-FL bond with PBP3-t/bocillin-FL bond. 50 µм protein was mixed with bocillin-FL with a molar ratio of 1:1 in PBS and incubated for one day at RT and subsequently transferred to 4 °C. The fluorescence polarization was measured after 5 days. **d**, Fluorescence polarization of PBP3-t/bocillin-FL complex over several days at 4 °C in PBS, with a 1:1 molar ratio of protein to bocillin-FL in PBS. Fluorescence polarization measurements consist of 3 separate replicates and the data indicate the mean ± SD.

As shown in Figure 1b, derivatives of ß-lactam antibiotics are expected to undergo ring opening when coupled to PenTag. This covalent conjugation could be a useful candidate for protein immobilization and purification, and moreover towards development of smart materials. To evaluate the functionality of PBP3-t compared to PBP3, we separately reacted both proteins with bocillin-FL penicillin, a fluorescent derivative of ß-lactam antibiotics, in a 1:1 protein to bocillin-FL molar ratio at room temperature. To compare the bond between bocillin-FL and both PBP3 versions, we measured fluorescence polarization of these conjugates (Figure 1c). No statistically significant difference could be seen between the polarization of PBP3 and PBP3-t.

Furthermore, we tested the stability of the bond between PBP3-t and bocillin-FL over time. We measured the fluorescence polarization of a mixture of PBP3-t and bocillin-FL over 7 days at 4 °C. We did not observe a significant difference in the polarization over time (Figure 1d). Moreover, we showed that our PenTag-antibiotic complex is most stable at physiological pH (between pH 7.5 and 8.5), which is of high relevance for biorthogonal conjugations (data shown in Supplementary Information, Figure S1b,c). A major challenge in protein overproduction in *E. coli* is the formation of insoluble aggregates.[42] One promising strategy to overcome this limitation is the fusion to solubilization domains[43], therefore, here we evaluated whether

PenTag can be used to increase solubility of an otherwise difficult-to-produce protein and showed its ability to enhance protein solubilization (data shown in Supplementary Information, **Figure** S2a).

### 2.2. PenTag as a novel synthesis strategy for stimuli-responsive hydrogels

After validating the interaction between PBP3 versions and ampicillin, we sought to extend this protein-ligand framework in biohybrid hydrogel synthesis. Programmable hydrogels are attractive materials for controlled delivery of biomolecules or as biosensing devices.[3,47,48] We aimed to develop stimuli-responsive hydrogels benefiting from the covalent bond between the PenTag and its small molecule ligand, ampicillin. For this, we first functionalized four-arm poly(ethylene glycol) succinimidyl carboxymethyl ester (4-arm PEG-SCM) with ampicillin (Supplementary Information **Figure** S3), followed by purification via reversed-phased preparative high-performance liquid chromatography (HPLC) to remove unbound ampicillin molecules (Supplementary Information **Figure** S4a). For the characterization of the 4-arm PEG-tetra ampicillin, we performed NMR and matrix-assisted laser desorption/ionization time-of-flight (MALDI ̶ ToF) of the purified samples (**Figure** S5, and Figure S4b respectively). The obtained results were in agreement with the expected NMR and MALDI ̶ ToF characterizations. As a crosslinker we designed an mCherry protein which was N- and C-terminally fused to PBP3-t via a linker containing cleavage sites for caspase-3 protease (Casp3) and analysed the possibility of the crosslinking within a single 4-arm PEG-ampicillin molecule (**Figure** S6). We then tested different ratios of protein to polymer (**Table** S2 and **Figure** S7) and monitored the mechanical behaviour of the two different stoichiometries (**Figure** S8). To synthesize stimuli-responsive hydrogels, we mixed 4-arm PEG-ampicillin with the protein crosslinker at PenTag : ampicillin-functionalized PEG-arm molar ratio of 1:4. Within 40 h, gelation was observed (**Figure** 2a). To test whether a controlled degradation of the hydrogel and release of mCherry is achievable by the stimulus, Casp3, we traced the fluorescence of mCherry in the supernatant after addition of different amounts of Casp3 (Figure 2b). Whereas no significant mCherry release was observed in the absence of Casp3, increasing protease concentrations correlated with higher mCherry release. These data show that PenTag offers a simple way of synthesizing biohybrid materials e.g., for the controlled, stimulus-dependent release of biomolecular cargos.

**Figure 2.**
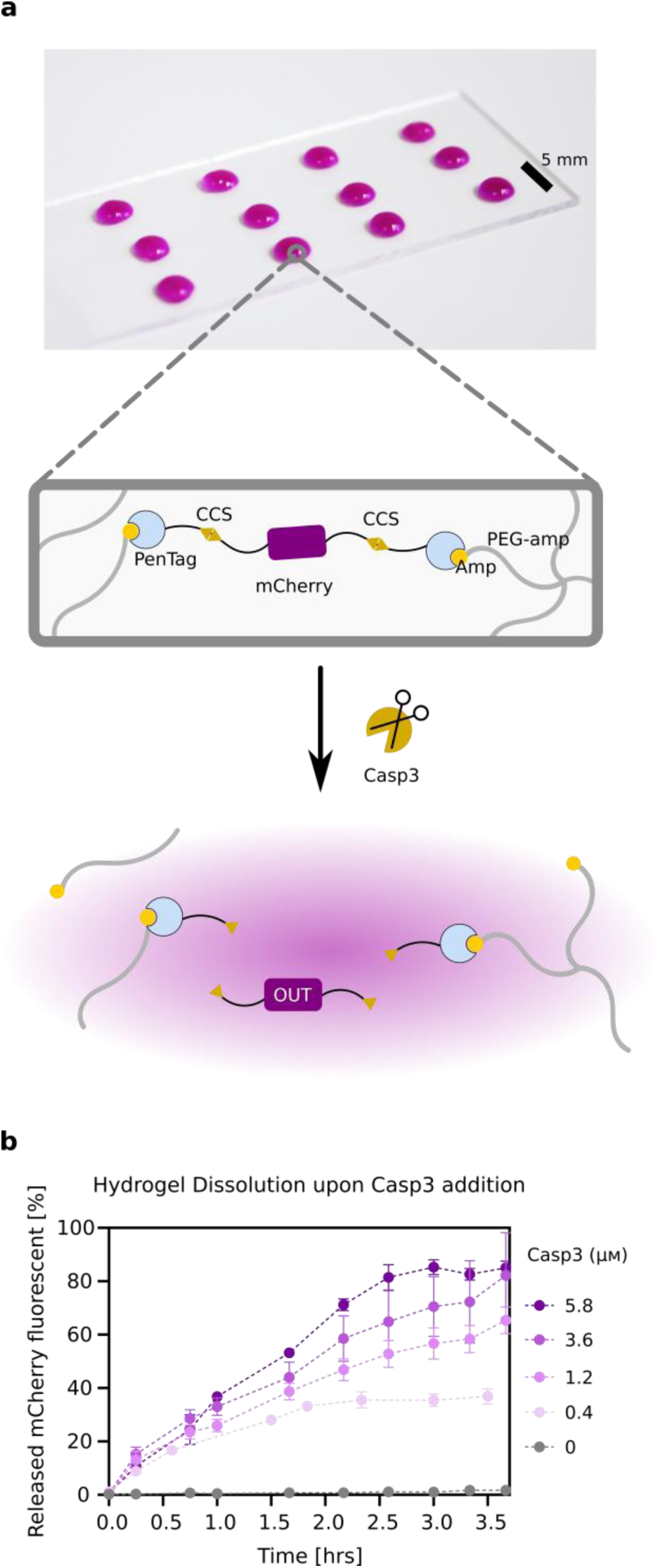
Synthesis of a stimuli-responsive hydrogel and its dissolution kinetics. **a**, Picture of synthesized hydrogels on a coated glass-slide and schematic of the building blocks involved in crosslinking. Hydrogels of 10 µl volume with PenTag : ampicillin-functionalized PEG-arm molar ratio of 1:4 and a final protein concentration of 70 mg ml^-1^ were synthesized and incubated for ∼40 h at RT. Subsequently, they were transferred to PBS for 6 h. **b**, Time-resolved dissolution of hydrogels upon addition of different Casp3 amounts in 200 µl PBS supplemented with 10 mм 2-β-mercaptoethanol (2-ME) was monitored by measuring the mCherry fluorescent of the supernatant at each time point. Maximum mCherry fluorescent intensity was measured after ∼5 h when complete gel dissolution was observed. Data represent 3-4 separate replicates for each condition, dotted lines connect the means of each condition and error bars show ±SD.

### 2.3. PenTag as affinity tag for immobilization of proteins on crosslinked agarose beads

Immobilization of proteins and biomolecules on solid supports enables the development of affinity matrices, micro devices, and bead-based protein assays for multiplexed analytical devices.[9,49,50] To produce an immobilization matrix to covalently bind PenTag, we functionalized cyanogen bromide (CNBr)-activated crosslinked agarose with ampicillin. CNBr first reacts with hydroxy groups of crosslinked agarose resulting in the formation of reactive cyanate ester groups. The primary amino group of ampicillin subsequently reacts with the cyanate ester, resulting in stable isourea linkages (**Figure** 3a).[51] To test whether ampicillin- functionalized beads can be used for the immobilization and subsequent release of PenTagged proteins, we genetically fused mCherry either to PBP3 or to PBP3-t. In both constructs, a protein linker containing the cleavage site for Casp3 was introduced to allow the release of mCherry from the beads by the addition of Casp3 (Figure 3b). To measure the binding capacity of the beads, we measured the fluorescence of Casp3-released mCherry and calculated the amount of released protein according to an mCherry standard per ml settled beads (Figure 3c). The binding capacity of >2 mg protein per ml settled ampicillin-functionalized beads compares favourably to the existing bead functionalization methods of HaloTag and SNAP-Tag with binding capacities of 7 mg and 1 mg protein per ml settled beads, respectively. To assess long-term stability of ampicillin-functionalized beads, the beads were lyophilized and reconstituted in buffer directly after lyophilization or after 18 days storage at 4°C. The binding capacity was assessed as described above and no statistically significant decline was observed in either condition. Moreover, no significant difference was observed between two PenTag versions, meaning that both of them can be used interchangeably. We further assessed PenTag as a purification tag for proteins and compared it to protein purification via IMAC and could show that PenTag can successfully be used as a purification tag as well. Data shown in Supplementary Information Figure S2b.

**Figure 3.**
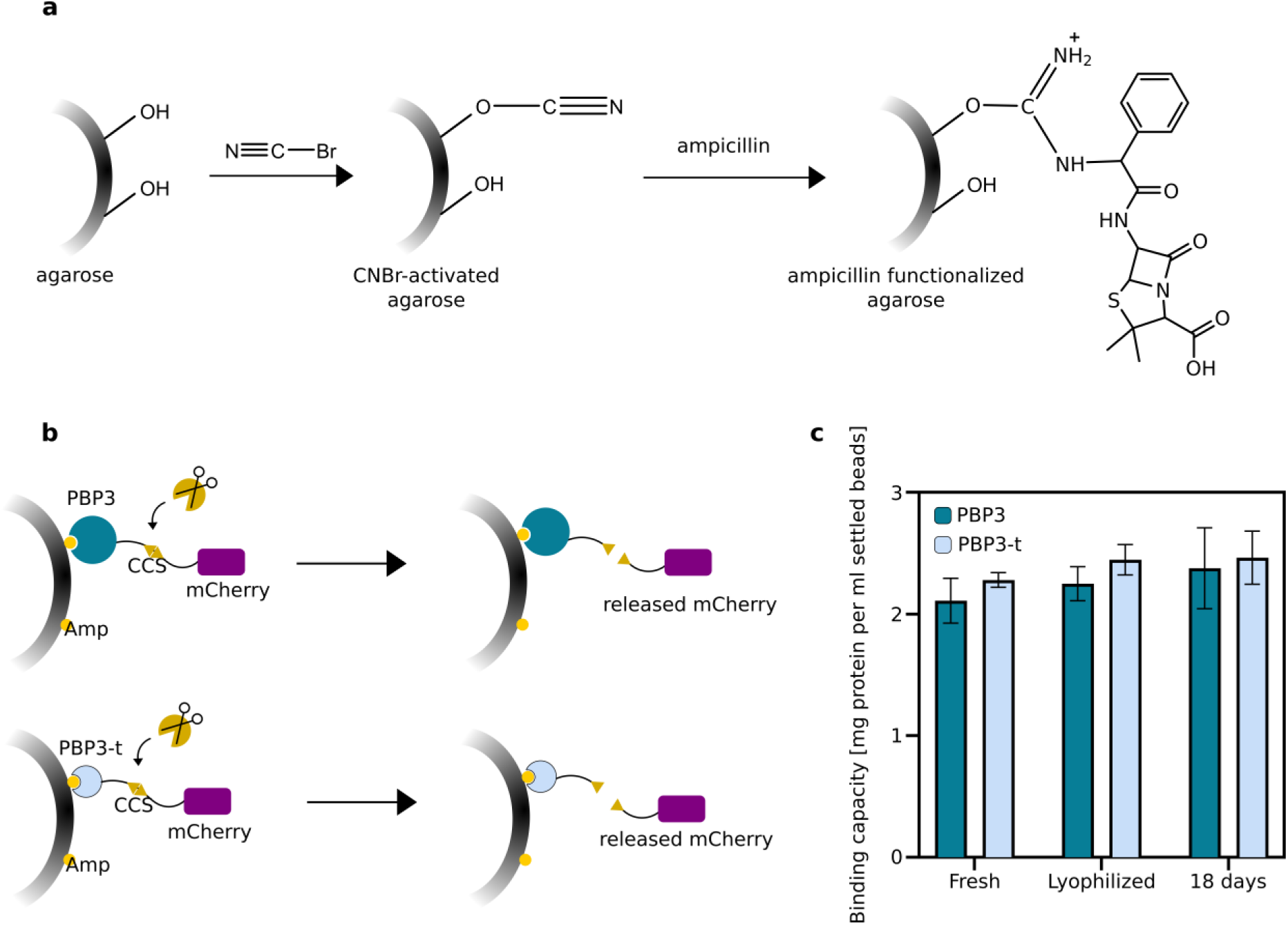
Conjugation of PenTagged proteins onto agarose beads. **a**, Functionalization of CNBr-activated agarose beads with ampicillin. **b**, Schematic representation of coupled PBP3-/or PBP3-t- fused mCherry on beads and mCherry release by adding Casp3 protease. **c**, mCherry release from ampicillin beads, captured by PenTag (either PBP3, or PBP3-t) at three different steps after functionalizing beads: right after functionalization, after lyophilization, and 18 days post-lyophilization. 80 µl of settled ampicillin-functionalized beads coupled with their respective proteins were transferred to 200 µl PBS supplemented with 10 mм 2-ME and 0.5 mg ml^-1^ Casp3 for one hour. The fluorescence of the supernatant was measured and correlated to the calibration of each protein. Binding capacity is reported as mg released protein per ml of settled beads. Bar charts represent the mean of 3 separate replicates and error bars indicate the ±SD.

### 2.4. Design, characterization, and development of an information processing material system

Biocomputing materials perceive several input stimuli, process those stimuli according to fundamental computational operations and produce a corresponding output. Examples include materials that can count the number of input pulses or that act as signal amplifier via integrated feedback and feedforward loops.[12,13,52] Materials with information-processing capacity enable multiple applications for example in diagnostics or personalized therapy. [16] Among computational operations, the encoder functionality is fundamental in binary data processing with manifold applications e.g. in therapies, medical image analysis, or biosensor fabrication. [53,54] The functionality and the operation characteristics of a 4:2 encoder as well as its implementation by connecting two OR gates is depicted in **Figure** 4a,b.

**Figure 4.**
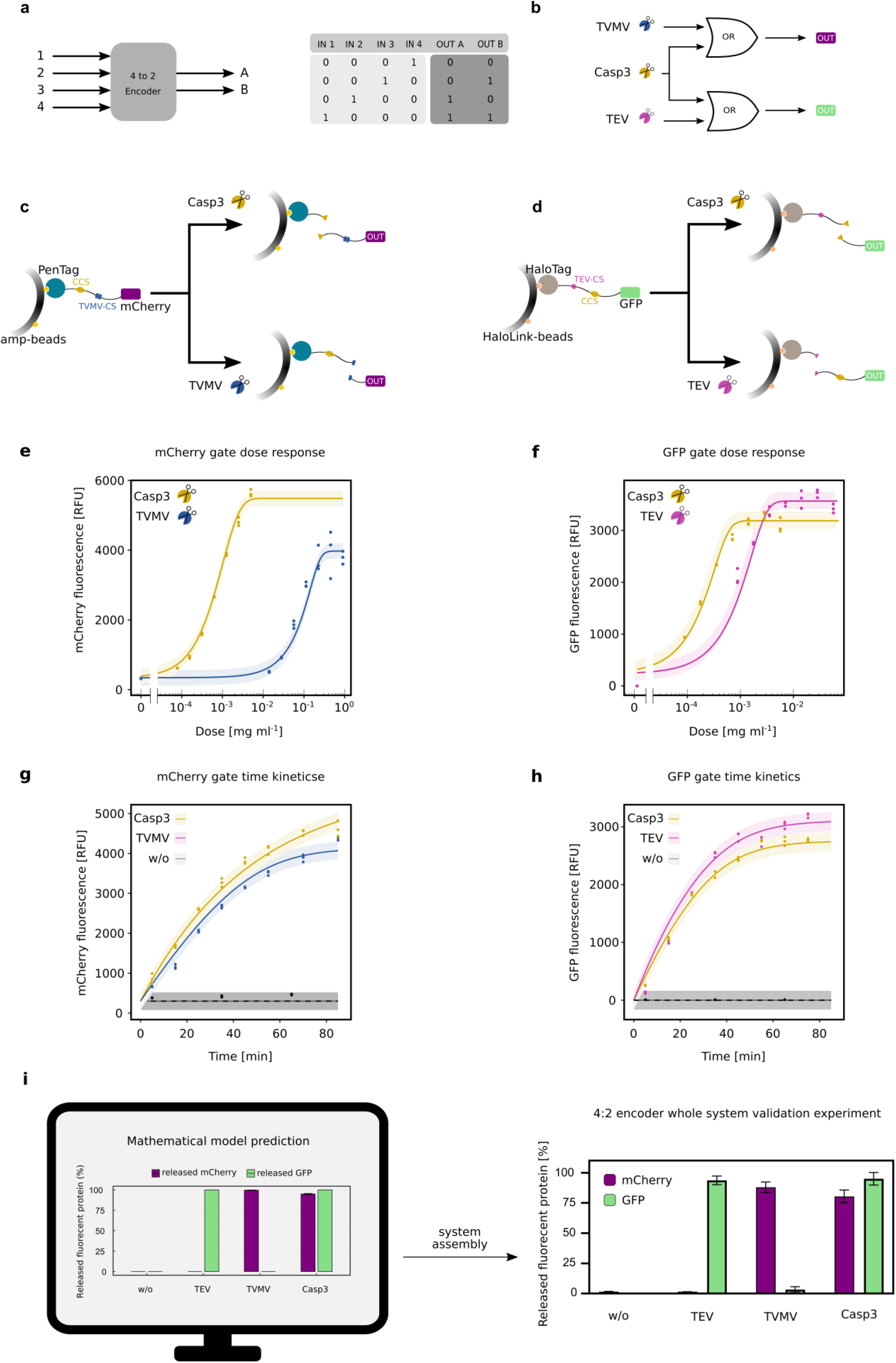
Development of two orthogonal material-based OR gates and their combination in a 4:2 encoder set-up. **a**, Schematic of a 4:2 encoder (left) and 4 different input/output combinations of such a 4:2 encoder (right). **b**, Combination of two different OR gates sharing one same input towards a 4:2 encoder. **c**, Schematic of mCherry OR gate. Output mCherry is supposed to be released in presence of either Casp3, or TVMV protease, or both. Molecular design of mCherry module consists of mCherry fused to PenTag by a linker containing cleavage sites for Casp3 and TVMV. Protein is bound through PenTag on ampicillin functionalized crosslinked agarose. **d**, Schematic of GFP OR gate. Output GFP is supposed to be released in presence of either Casp3, or TEV protease, or both. Molecular design of GFP module, consisting of GFP fused to HaloTag by a linker containing cleavage sites for TEV and Casp3. Output GFP is released in presence of each of proteases. Protein is bound through HaloTag on HaloLink material. **e**, Data and model fits of the dose-dependent release of mCherry from ampicillin material module upon addition of different concentrations of TVMV and Casp3. 7 µl settled beads coupled with mCherry protein were transferred to 200 µl assay buffer (PBS, 10 mм 2-ME, 0.001% (v/v) Tween-20) and supplemented with the respective protease concentrations. The mCherry fluorescence of the supernatant was measured after 1 h. **f**, Data and model fits of the dose-dependent release of the GFP from HaloLink material module upon addition of different concentrations of Casp3 and TEV. 5 µl settled beads coupled with GFP protein were transferred to 200 µl assay buffer and supplemented with the respective protease concentrations. The GFP fluorescence of the supernatant was measured after 1 h **g**, Data and model fits of the time-resolved measurements of mCherry module upon addition of Casp3 or TVMV over time. Reactions containing 7 µl settled beads in the assay buffer were supplemented with the respective proteases, 1.25 μg ml^-1^ Casp3 and 225 μg ml^-1^ for TVMV. Samples from the supernatant were taken and measured at each time point. h, Data and model fits of the time-resolved measurements of GFP module upon addition of TEV or Casp3 over time. Reactions containing 5 µl settled beads in the assay buffer were supplemented with the respective proteases, 0.625 μg ml^-1^ Casp3 and 3.125 μg ml^-1^ for TEV. Samples from the supernatant were taken and measured at each time point. **i**, Combination of both systems towards a 4:2 encoder system predicted by the mathematical model (left) and validated by experiments (right) with the concentrations predicted by the model: 6, 14, and 800 μg ml^-1^ for Casp3, TEV, and TVMV respectively. Maximum RFU values were measured after 55 min and data is normalized to this value. Bar chart indicates the mean ± SD. Samples were prepared and measured as triplicates, and for negative controls duplicates were measured.

To assemble an encoder material framework, we capitalized on our ampicillin-functionalized agarose beads for protein immobilization via PenTag. We first developed two separate agarose- based orthogonal OR gates (Figure 4c,d), each being responsive to two different proteases and releasing a fluorescent protein, mCherry or GFP, as output. The first OR gate is composed of mCherry genetically fused to PenTag via a linker containing cleavage sites for Casp3 as well as for the tobacco vein mottling virus (TVMV) protease. The fusion protein is covalently coupled to ampicillin-functionalized crosslinked agarose. The addition of either protease is expected to result in the release of mCherry that can be quantified by fluorescence measurement (Figure 4c). The second OR gate is synthesized from GFP protein genetically fused to HaloTag via a linker containing cleavage sites for tobacco etch virus protease (TEV) and Casp3. This fusion protein is covalently coupled to Halo-ligand-functionalized crosslinked agarose, commercially available HaloLink resin. The addition of TEV OR Casp3 results in the release of GFP that is quantified by fluorescence measurement (Figure 4d).

To characterize each of the OR gates, we first monitored dose-dependent release of mCherry and GFP upon addition of their input proteases (Figure 4e and Figure 4f, respectively). We further characterized the cleavage kinetics in response to protease addition. For this, we fixed the protease concentration to the lowest value that resulted in a plateau in the above dose- response measurements (for the mCherry gate: 1.25 µg ml^-1^ for Casp3 and 225 µg ml^-1^ for TVMV; for the GFP gate: 0.625 µg ml^-1^ for Casp3 and 3.125 µg ml^-1^ for TEV). The kinetic data from these experiments were used to parameterize a quantitative mathematical model based on ordinary differential equations (ODEs). In order to identify a parameter combination in which the assembled system would show 4:2 encoder functionality within a desired timeframe (30 min), we simulated the performance of the assembled system as a function of the concentrations of all input proteases (Figure 4i left) (details of the mathematical model can be found in Supplementary Information, **Figure** S9, S10 and **Table** S3).

To confirm the model predictions, an experiment was conducted with the following predicted parameters: final concentrations of 6, 14, and 800 µg ml^-1^ for Casp3, TEV, and TVMV respectively. To assemble the 4:2 encoder, we combined both modules with each other in a single pot in the presence or absence of each of the three input proteases (time-resolved model predictions for the encoder response are shown in **Figure** S11a). An output pattern specific for a 4:2 encoder was predicted (Figure 4i left). This pattern could experimentally be obtained, which demonstrates the functionality of the overall computing network and validates the predictions of the mathematical model (Figure 4i right, kinetic data in Supplementary Information Figure S11b).

This set-up shows that this novel conjugation platform is biorthogonal to existing protein tags (HaloTag) and can further serve as a robust building block towards the development of information-processing materials.

## 3. Discussion

We developed a truncated version of PBP3 and demonstrated the covalent binding of both PBP3 versions to ampicillin and termed this technology as PenTag. The spontaneous binding of PenTag to its small molecule ligand opened the door for multiple applications in chemical biology and biorthogonal coupling. Protein tags that not only solve purification issues of proteins of interest, but also facilitate protein labelling by organic dyes are extremely beneficial for protein visualization and detection.[32] An overview of some current protein-tagging systems compared to PenTag can be found in Table S4. Moreover, embedding proteins of interest into hydrogels through a covalent protein-ligand interaction can serve as attractive means for the immobilization and controlled delivery of proteins and other molecules.[9,16] Furthermore, such a system that is orthogonal to the existing systems, can extend the library for covalent protein immobilization on materials and enable the development of information-processing materials.[10] A drawback of the already existing systems is the expensive and difficult-to-synthesize small molecules. Here, we benefit from ampicillin as the ligand, a small, available, and cheap molecule that can easily be coupled via its primary amino group to a large variety of N-hydroxy succinimide (NHS)-functionalized commercially available reagents. For example, NHS-functionalized dyes, biotin-labelling reagents, surfaces, polymers, and beads are readily available.[37,55–59] However, it has to be noted that upon coupling of ampicillin to PenTag, the ß-lactam ring is opened and thereby the antibiotic activity of ampicillin is abolished.

In order to introduce a multi-functional protein tag, we first engineered a truncated version of PBP3 with a size of 30.2 kDa, which is 12 kDa smaller than the previous PBP3 version, yet as active as the 42 kDa version. PenTag was shown to enhance the solubility of a fused insoluble protein of interest, to address a common challenge when working with proteins. [60] Our system has shown to be stable at physiological pH values, the environment where most proteins are stable at. Despite the increase in size when fusing PenTag to proteins of interest, this method offers high specificity and does not require addition of an enzyme as for enzyme-based conjugation methods.[61–63] This facilitates downstream processing of functionalized materials. We used PenTag strategy for covalent immobilization of proteins of interest into hydrogels and their controlled release by defined environmental cues. Due to the easy coupling of ampicillin to other molecules, different polymers with various sizes can be functionalized with ampicillin, and thereby desired hydrogels can be tailored depending on the final applications. Here, we chose a 4-arm PEG-SCM and functionalized it with ampicillin and could establish covalent crosslinks using PenTag where a controlled release of protein of interest from the hydrogel was achieved by addition of Casp3 protease. Due to the cost-efficiency of this conjugation technique, this tool can be used for higher-scale applications for example in affinity chromatography for the industrial purification of proteins, or in wound dressings for patients with major burn injuries, or in water-treatment for example towards water desalination, removal of toxic impurities like heavy metals, organic dyes, etc.[64–66]

Furthermore, we synthesized ampicillin functionalized crosslinked agarose that has applications in various fields, such as protein microdevice assays or for high purity purifications.[49,67] Lyophilized ampicillin agarose beads were shown to be stable, which allows the off-the-shelf usage for protein immobilization or capture.[68]

Signalling and logic circuit designs using proteases have significantly been advanced in the recent years.[13,69,70] Implementation of such circuits onto materials frameworks would add another level of applicability and dimension towards development of powerful information-processing materials.[11,12] Benefiting from various synthetic biological building blocks, we integrated conceptual engineering principles into materials. These materials sense environmental cues as their inputs, can then process information, and finally exhibit a regulated output. In this work, we first assembled simple OR gate material modules using PenTag and compared it to the existing systems. We then did thorough characterization of the both systems by a mathematical model, and finally developed a one-pot combined system that could perform more complicated information processing and produce regulated and clear outputs in a 4:2 encoder design manner. We further validated the predictions of the mathematical model by experimental data. The concepts and building blocks introduced here, lay the foundation for the development of various information-processing materials for advanced biomedical applications. For example, an anti-cancer antibody such as Bevacizumab can N-/ and C-terminally be fused to PenTags and anchored onto a material support via linkers that contain the cleavage sites for cancer-associated proteases, for example matrix metalloproteases (MMPs) or cathepsins in metastatic cancer progressions.[71–73] This would result in an AND-Gate set-up that releases the anticancer antibody in response to the presence of both cancer-associated proteases.

## 4. Conclusion

In summary, we introduced a specific and multifunctional protein tag which is highly modular regarding the protein of interest, and even useful when working with insoluble proteins of interest. We showed broad applications for PenTag and specifically demonstrated that hydrogels and other materials systems can be tailored to response to environmental cues, or to perform sensory functions using this strategy.

We see PenTag as a versatile, modular tool, orthogonal to existing systems, which is an advantageous candidate for implementation and detection of biological molecules in material frameworks by means of a simple small molecule. Due to its simplicity, cost-efficiency, and high specificity, proteins of interest could covalently and stably be coupled onto different substrates such as hydrogels and other solid supports. We expect that this platform will find broad applications not only in synthetic biology and protein-based biohybrid materials, but also in advanced devices for analytical and diagnostic applications, as controlled drug delivery vehicles, or in tissue engineering and regenerative medicine.

## 5. Experimental Section/Methods

### Cloning of the plasmids

The amino acid sequence of all plasmids and their description are listed in Supplementary Information Table S1. Constructs were cloned by Gibson assembly[74].

### Protein expression and purification

For production of all recombinant proteins, plasmids were transformed into *E. coli* BL21 (DE3) pLysS (Invitrogen). Cells were grown at 37 °C while shaking at 150 rpm in LB medium containing 100 µg ml^-1^ ampicillin (Carl Roth) and 36 µg ml^-1^ chloramphenicol (Carl Roth). Once an OD600 of 0.6 was reached, the cultures were induced with 1 mм isopropyl ß-D-1- thiogalactopyranoside (IPTG) and incubated overnight at 18 °C or for 4 h at 30 °C or 37 °C (pKJ57 4 h at 37 °C). Cells were harvested by centrifugation at 6000 x *g* for 10 min and were subsequently resuspended in 35 ml nickel-lysis buffer (50 mм NaH2PO4, 300 mм NaCl, 10 mм imidazole, pH 8.0), flash frozen and stored at -80 °C until further use. For lysis, cells were sonicated at 60% amplitude and 0.5 s/ 1 s pulse/ pause intervals with a Sonopuls HD3100 (Bandelin). Crude lysates were centrifuged at 30,000 x *g* for 30 min. The protein in the supernatant were purified IMAC, using Ni-NTA Superflow agarose beads (Qiagen) or using ampicillin functionalized agarose. For Ni-NTA purification following the manufacturer’s instructions, proteins were loaded onto the already Ni lysis-equilibrated columns, unbound protein was washed away by two wash steps using Ni wash buffer (50 mм NaH2PO4, 300 mм NaCl, 20 mм Imidazole, pH 8.0), followed by elution with Ni elution buffer (50 mм NaH2PO4, 300 mм NaCl, 250 mм Imidazole, pH 8.0). Eluted proteases were additionally supplemented with 10 mм 2-ME. All eluted proteins were dialyzed against 5 L 1x PBS using SnakeSkin dialysis tubes with 3.5k molecular weight cut-off (MWCO) (Thermo Fisher Scientific) at 4 °C. Proteins were subsequently analysed by SDS-PAGE (15% (w/v) gels at 160 V), stained by Coomassie brilliant blue staining, and the concentration of the protein was determined either by SDS-PAGE or by Bradford assay using protein assay dye reagent (Bio-Rad) and bovine serum albumin (Carl Roth) as standard. Measurement was performed in a Multiskan GO spectrophotometer at 575 nm (Thermo Fisher Scientific). In some cases, proteins were concentrated using Spin-X UF 6 columns (Corning Inc., Corning, USA). Purified proteins were supplemented with 15% (v/v) glycerol and stored at -80 °C for further use.

For solubility tag experiments, PBP3-t was N-terminally fused to an insoluble construct containing a Pyruvate dehydrogenase complex repressor (PdhR) and an intrinsically disordered protein domain. Protein expression was performed at 18 °C for 16 h.

### Fluorescence polarization

To analyze the binding between ampicillin and PenTag, a fluorescent derivate of penicillin, Bocillin-FL (Thermo Fisher Scientific), was incubated with both PBP3 versions in a molar ratio of 1:1 at 4 °C, with a final concentration of 50 µм in 1x PBS. Blank samples in the absence of protein were prepared to measure background fluorescence and the fluorescence quenching of the fluorophore when incubated for several days at 4 °C. The mixtures were allowed to reach equilibrium for 1 h at room temperature and then transferred to 4 °C. Fluorescence polarization of 15 µl of the samples in a 384-well plates (Corning 3575) was measured by exciting the samples with polarized light at 485/20 nm and emission at 528/20 nm at a parallel and perpendicular angle for 40 min with an interval of 5 min to make sure the plateau is maintained using a Synergy 4 plate reader (BioTek Synergy 4, Vermont, United States). Fluorescence polarization was then calculated using the following equation:

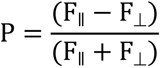

Fǁ = fluorescence parallel to the excitation plane; F⊥ = fluorescence perpendicular to the excitation plane). As negative control the FP of Bocillin-FL alone was measured and its fluorescence quenching over days was used for data normalization.

### Functionalization of 4-arm PEG with ampicillin

4-arm PEG-NHS, MW 5k (4-arm PEG-SCM, Creative PEG works, Durham, USA) and ampicillin (Roth) were both dissolved in dimethylformamide (DMF) with a concentration of 100 mg ml^-1^ and 20 mg ml^-1^ respectively. A 6x molar excess of ampicillin (192 µmol) was mixed with 15 µmol triethylamine (Sigma) prior to being added drop-wise to PEG-SCM (32 µmol). All steps were conducted in an exicator under argon flow. For purification, preparative HPLC (Agilent Technologies 1260 Infinity I/II) was conducted to separate excess ampicillin from PEG-tetra-ampicillin. Briefly, preparative HPLC was employed using a reversed-phase C18 column (Macherey-Nagel VP 250/21 Nucleodur 100-5) with a mobile phase solvent A (Water + 0.05% vol/vol trifluoroacetic acid (TFA)), solvent B (acetonitrile + 0.05% vol/vol TFA). The column was eluted with a gradient of solvent B 10-90% over 20 minutes at a flow rate of 20 ml min^-1^, and fractions were monitored by UV-absorption at 230 nm.

### Characterization of the polymer

Samples of collected fractions of preparative HPLC were monitored by MALDI-ToF mass spectrometry to confirm the right size of the purified samples. For this, samples were dissolved in acetonitrile (HPLC grade). 5 µl of the sample solution was mixed with 5 µl of a saturated solution of α-cyano-4-hydroxycinnamic acid matrix for MALDI-MS (Sigma-Aldrich) in acetonitrile. 1.5 µl of the latter mixture was applied to the MALDI target and let dry. Analysis was carried out on a Microflex MALDI mass spectrometer (Bruker) and data was processed via mMass software and GraphPad Prism.

For precise characterization of the final product and free ampicillin molecule, ^1^H NMR spectra were recorded at room temperature on a Bruker Avance Neo (400 MHz). Samples were dissolved in deuterated DMSO or chloroform.

Once the fractions were characterized, they were freeze-dried and lyophilized. Samples were subsequently diluted in ddH2O and stored at -20 °C for further use. The concentration of the PEG solution was determined by UV-Vis according to a calibration standard of ampicillin molecule (0 – 10 mg ml^-1^) to measure the PEG-coupled ampicillin moieties.

### Hydrogel synthesis

For hydrogel synthesis, purified PBP3t-mCherry-PBP3t (pHM528) was concentrated to ∼150 mg ml^-1^ by centrifugation using Spin-X UF 20 ml centrifugal concentrators (30 kDa MWCO, Corning Inc., Corning, USA). Hydrogels of 10 µl volume were produced by 70 mg ml^-1^ crosslinker protein (pHM528) with PEG-tetra-ampicillin in PBS in PenTag : ampicillin- functionalized PEG-arm molar ratio of 1:4 . Glass slides were siliconized by Sigmacote (Sigma-Aldrich) according to manufacturer’s instructions. Briefly, clean glass slides were immersed in Sigmacote, air-dried under a fume hood, and were subsequently washed with water and let dry. The polymer-protein mixture was spotted on siliconized glass slides and incubated at room temperature for ∼40 h under humidifying atmosphere. Afterwards, the hydrogels were transferred to 1 ml 1x PBS and incubated for 6 h to let the hydrogels swell. Swollen hydrogels were subsequently transferred to different experimental set-ups and incubated for 2 h in 200 µl 1x PBS with 10 mм 2-ME and at room temperature on an orbital shaker. Fluorescent measurements of the supernatant were carried out at desired time points at 575/620 nm Ex/Em and the % dissolution was measured by dividing the fluorescent intensity by the maximum mCherry fluorescent intensity upon complete gel dissolution, after incubation with 3.6 and 5.8 µм Casp3 for 285 min

### Functionalization of CNBr-activated Sepharose 4B beads

Ampicillin-beads were produced by coupling ampicillin to CNBr-activated Sepharose beads (GE Healthcare). Coupling was performed according to manufacturer’s instructions. Briefly, 1 g of the lyophilized beads were washed for 30 min with ice cold 1 mм HCl to preserve the activity of the reactive groups. The beads were then equilibrated with coupling buffer (0.1 м NaHCO3, 0.5 м NaCl, pH 8.3) and subsequently 1 ml ampicillin (4.57 mм diluted in water) was added to the beads. The mixture was incubated overnight at 4 °C on a rotary shaker. The next day, excess ampicillin was washed away with coupling buffer. To block any remaining active groups, the beads were slowly agitated for 2 h in blocking buffer (0.1 М Tris-HCl, pH 8.0) followed by four wash cycles with alternating pH buffers (0.1 м acetic acid/sodium acetate with 0.5 м NaCl, pH 4 and 0.1 м Tris-HCl with 0.5 м NaCl, pH 8.0). Finally, beads were washed with deionized water (ddH2O) and transferred to 10 mg ml^-1^ BSA dissolved in water and incubated overnight at 4 °C on a rotary shaker. After coupling, the beads were lyophilized (Christ Alpha 1-4 LSCbasic, Germany) and stored at 4 °C for further use.

### Binding capacity and purification tag

For binding capacity tests, 500 µl of the settled ampicillin beads were incubated overnight at 4°C with 4.5 mg of protein PBP3-CCS-mCherry (pHM151) or PBP3t-CCS-mCherry (pHM525). On the following day, the beads were washed four times with 1x PBS and the supernatant of the wash fractions was kept for later measurements. To measure the binding capacity, 80 µl of the settled beads bound to the protein were added to 200 µl 1x PBS containing 10 mм 2-ME and 20 µl of Casp3 (0.5 mg ml^-1^). After incubating for 1 h the relative fluorescence unit (RFU) of the supernatant was measured at 575/620 nm Ex/Em using an Infinite M200 pro micro plate reader (Tecan). The fluorescence was correlated to dilution series of each protein with concentrations between 0 and 1.5 mg ml^-1^ and thereby the binding capacity was reported as mg protein per ml settled beads.

### Encoder system

Prior to assembling the encoder system, each OR-gate was established separately. OR gate 1 was prepared by immobilizing PBP3-fused mCherry protein (pHM151) onto ampicillin- functionalized crosslinked material as explained in “binding capacity” section. To characterize this system, a dose-dependent mCherry release was monitored by the addition of different concentrations of TEV (pRK793) and Casp3 (pKJ57) proteases, using 7 µl settled beads per reaction, all reactions in 200 µl 1x PBS supplemented with 10 mм 2-ME and 0.001 % (v/v) Tween-20 and the respective protease concentrations. For time kinetic measurements, a specific concentration for each protease was chosen (1.25 µg ml^-1^ for Casp3 and 225 µg ml^-1^ for TVMV).

For each time point, a 200 µl reaction mixtures was prepared. Samples were taken in triplicates and measured at 575/625 nm Ex/Em as explained above. For OR Gate 2, HaloTagged GFP (pHM150) was coupled onto HaloLink™ resin (Promega) following the manufacturer’s instructions. Briefly, 500 µl of settled beads were equilibrated with binding buffer (100 mм Tris/HCl, 150 mм NaCl, 0.5% Triton, pH 7.6) and incubated with 3.6 mg GFP protein on a rotary shaker overnight at 4 °C. On the following day, beads were washed 4 times with binding buffer. Conduction of the dose-response and kinetic measurements were similar to the first OR gate. For dose response measurements, the only difference was that 5 µl of settled beads were used for each condition. For time kinetic measurements, 0.625 µg ml^-1^ Casp3 and 3.125 µg ml^-1^ TEV were used. For the encoder system, both proteins were coupled onto their respective material frameworks and the crosslinked materials (5 µl GFP settled beads, and 7 µl mCherry settled beads) were mixed together in 2 ml 1x PBS, supplemented with 10 mм 2-ME and final concentrations of 6, 14, and 800 µg ml^-1^ for Casp3, TEV, and TVMV were added respectively. The measured RFU for each gate was normalized to the maximum RFU values after 55 min and values were reported as percentage of released proteins. A time kinetic measurement was performed for the encoder system combination for 55 min every 5 min.

### Statistical analysis

Mean values of at least three separate biological replicates were shown with error bars representing ± SD. Statistical difference was assessed by ordinary one-way ANOVA or unpaired t-test (Figure 1 c) using GraphPad Prism 9.2.0 and assessed as not significant where reported in the manuscript. Normalizations are reported in each respective section.

## Supporting information

Supplementary Tables 1-4, Supplementary Figures 1-11 and supplementary method for mathematical model

## Supporting Information

Supporting Information is available from the Wiley Online Library or from the author.

## Acknowledgements

This work was supported by the European Union (ERC, STEADY, 101053857), the German Research Foundation (Deutsche Forschungsgemeinschaft, DFG) under Germany’s Excellence Strategy –CIBSS, EXC-2189, Project ID: 390939984 and under the Excellence Initiative of the German Federal and State Governments – BIOSS, EXC-294, and in part by the Ministry for Science, Research and Arts of the state of Baden-Württemberg. The authors acknowledge the support of the Freiburg Galaxy Team funded by Collaborative Research Centre 992 Medical Epigenetics (DFG grant SFB 992/1 2012) and the German Federal Ministry of Education and Research (BMBF grant 031 A538A de.NBI-RBC).

## Data Availability Statement

The data that supports the findings of this study are available in the supplementary material of this article

## Supporting Information

Supporting Information is available from the Wiley Online Library or from the author.

Received: ((will be filled in by the editorial staff)) Revised: ((will be filled in by the editorial staff)) Published online: ((will be filled in by the editorial staff))

