## Supplementary Tables 1-4, Supplementary Figures 1-11 and supplementary method for mathematical model for "PenTag, a Versatile Platform for Synthesizing Protein-Polymer Biohybrid Materials"

### Contents

Supplementary tables 1-4

Supplementary figures 1-11

Supplementary method: description of the mathematical model

**Table S1. List of constructs.**

| Plasmid | Amino acid sequence | Description | Reference |
|---|---|---|---|
| **pHM69** | MKIEEGKLVIWINGDKGYNGLAEVGKKFEKDTGIKVTVEHPDK<br>LEEKFPQVAATGDGPDIIFWAHDRFGGYAQSGLLAEITPDKAFQ<br>DKLYPFTWDAVRYNGKLIAYPIAVEALSLIYNKDLLPNPPKTWE<br>EIPALDKELKAKGKSALMFNLQEPYFTWPLIAADGGYAFKYEN<br>GKYDIKDVGVDNAGAKAGLTFLVDLIKNKHMNADTDYSIAEA<br>AFNKGETAMTINGPWAWSNIDTSKVNYGVTVLPTFKGQPSKPF<br>VGVLSAGINAASPNKELAKEFLENYLLTDEGLEAVNKDKPLGA<br>VALKSYEEELAKDPRIAATMENAQKGEIMPNIPQMSAFWYAVR<br>TAVINAASGRQTVDEALKDAQTNSSSNNNNNNNNGGGSGGGS<br>GGGSSKALLKGVRDFNPISACVCLLENSSDGHSERLFGIGFGPYI<br>IANQHLFRRNNGELTIKTMHGEFKVKNSTQLQMKPVEGRDIIVI<br>KMAKDFPPFPQKLKFRQPTIKDRVCMVSTNFQQKSVSSLVSESS<br>HIVHKEDTSFWQHWITTKDGQCGSPLVSIIDGNILGIHSLTHTTN<br>GSNYFVEFPEKFVATYLDAADGWCKNWKFNADKISWGSFTLV<br>EDAPEDHHHHHH<br>MalE/TVMV/HisTag | Sequence coding for Maltose binding protein fused to TVMV protease and a His-Tag for purification (used in encoder) | This work |
| **pHM150** | MEIGTGFPFDPHYVEVLGERMHYVDVGPRDGTPVLFLHGNPTS<br>SYVWRNIIPHVAPTHRCIAPDLIGMGKSDKPDLGYFFDDHVRFM<br>DAFIEALGLEEVVLVIHDWGSALGFHWAKRNPERVKGIAFMEFI<br>RPIPTWDEWPEFARETFQAFRTTDVGRKLIIDQNVFIEGTLPMG<br>VVRPLTEVEMDHYREPFLNPVDREPLWRFPNELPIAGEPANIVA<br>LVEEYMDWLHQSPVPKLLFWGTPGVLIPPAEAARLAKSLPNCK<br>AVDIGPGLNLLQEDNPDLIGSEIARWLSTLEISGHGGGSGGGSG<br>GENLYFQSGGGSGGGDEVDGGGSGGGSGGMVSKGEELFTGVV<br>PILVELDGDVNGHKFSVSGEGEGDATYGKLTLKFICTTGKLPVP<br>WPTLVTTLTYGVQCFSRYPDHMKQHDFFKSAMPEGYVQERTIF<br>FKDDGNYKTRAEVKFEGDTLVNRIELKGIDFKEDGNILGHKLE<br>YNYNSHNVYIMADKQKNGIKVNFKIRHNIEDGSVQLADHYQQ<br>NTPIGDGPVLLPDNHYLSTQSKLSKDPNEKRDHMVLLEFVTAA<br>GITLGMDELYKGGSGGHHHHHH<br>Halotag/TEVS/CCS/EGFP/HisTag | Sequence coding for HaloTag fused to enhanced green fluorescent protein via a linker containing a TEV and Casp3 protease cleavage site and a HisTag for purification (used in encoder set-up as GFP gate) | This work |
| **PHM151** | MGVSTAVAQDFTIAAKHAIAVEANTGKILYEKDATQPVEIASIT<br>KLITVYLVYEALENGSITLSTPVDISDYPYQLTTNSEASNIPMEA<br>RNYTVEELLEATLVSSANSAAIALAEKIAGSEKDFVDMMRAKL<br>LEWGIQDATVVNTTGLNNETLGDNIYPGSKKDEENKLSAYDVA<br>IVARNLIKKYPQVLEITKKPSSTFAGMTITSTNYMLEGMPAYRG<br>GFDGLKTGTTDKAGESFVGTTVEKGMRVITVVLNADHQDNNP<br>YARFTATSSLMDYISSTFTLRKIVQQGDAYQDSKAPVQDGKED<br>TVIAVAPEDIYLIERVGNQSSQSVQFTPDSKAIPAPLEAGTVVGH<br>LTYEDKDLIGQGYITTERPSFEMVADKKIEKGSGGGSGGDEVD<br>GSGGGSGGETVRFQSGSGGGSGGAIIKEFMRFKVHMEGSVNGH<br>EFEIEGEGEGRPYEGTQTAKLKVTKGGPLPFAWDILSPQFMYGS<br>KAYVKHPADIPDYLKLSFPEGFKWERVMNFEDGGVVTVTQDSS<br>LQDGEFIYKVKLRGTNFPSDGPVMQKKTMGWEASSERMYPED | Sequence coding for PBP3 coupled to c-terminally His-tagged mCherry fluorescent protein via a linker containing Casp3 protease and TVMV protease cleavage sites (used for beads and in encoder set- | This work |

| | | | |
|---|---|---|---|
| | GALKGEIKQRLKLKDGGHYDAEVKTTYKAKKPVQLPGAYNVNIKLDITSHNEDYTIVEQYERAEGRHSTGGMDELYKGGSGGHHHHHH<br>PBP3/CCS/TVMVS/mCherry/HisTag | up as mCherry gate) | |
| **pHM512** | MGVSTAVAQDFTIAAKHAIAVEANTGKILYEKDATQPVEIASITKLITVYLVYEALENGSITLSTPVDISDYPYQLTTNSEASNIPMEARNYTVEELLEATLVSSANSAAIALAEKIAGSEKDFVDMMRAKLLEWGIQDATVVNTTGLNNETLGDNIYPGSKKDEENKLSAYDVAIVARNLIKKYPQVLEITKKPSSTFAGMTITSTNYMLEGMPAYRGGFDGLKTGTTDKAGESFVGTTVEKGMRVITVVLNADHQDNNPYARFTATSSLMDYISSTFHHHHHH<br>PenTag/HisTag | Truncated version of PBP3 (PBP3-t) | This work |
| **pHM525** | MGVSTAVAQDFTIAAKHAIAVEANTGKILYEKDATQPVEIASITKLITVYLVYEALENGSITLSTPVDISDYPYQLTTNSEASNIPMEARNYTVEELLEATLVSSANSAAIALAEKIAGSEKDFVDMMRAKLLEWGIQDATVVNTTGLNNETLGDNIYPGSKKDEENKLSAYDVAIVARNLIKKYPQVLEITKKPSSTFAGMTITSTNYMLEGMPAYRGGFDGLKTGTTDKAGESFVGTTVEKGMRVITVVLNADHQDNNPYARFTATSSLMDYISSTFGGGSGGGSDEVDGGGPAGEASSIPNREGKPIPNPLLGLGSTRTGEVSKGEEDNMAIIKEFMRFKVHMEGSVNGHEFEIEGEGEGRPYEGTQTAKLKVTKGGPLPFAWDILSPQFMYGSKAYVKHPADIPDYLKLSFPEGFKWERVMNFEDGGVVTVTQDSSLQDGEFIYKVKLRGTNFPSDGPVMQKKTMGWEASSERMYPEDGALKGEIKQRLKLKDGGHYDAEVKTTYKAKKPVQLPGAYNVNIKLDITSHNEDYTIVEQYERAEGRHSTGGMDELYKHHHHHH<br>PenTag/CCS/mCherry/HisTag | Sequence coding for a truncated penicillin binding protein 3 coupled to a C-terminal mCherry via a linker containing a caspase-3 cleavage site with a C-terminal His-Tag (for binding capacity test) | This work |
| **pHM528** | MGVSTAVAQDFTIAAKHAIAVEANTGKILYEKDATQPVEIASITKLITVYLVYEALENGSITLSTPVDISDYPYQLTTNSEASNIPMEARNYTVEELLEATLVSSANSAAIALAEKIAGSEKDFVDMMRAKLLEWGIQDATVVNTTGLNNETLGDNIYPGSKKDEENKLSAYDVAIVARNLIKKYPQVLEITKKPSSTFAGMTITSTNYMLEGMPAYRGGFDGLKTGTTDKAGESFVGTTVEKGMRVITVVLNADHQDNNPYARFTATSSLMDYISSTFGGGSGGGSDEVDGGGPAGEASSIPNREGKPIPNPLLGLGSTRTGEVSKGEEDNMAIIKEFMRFKVHMEGSVNGHEFEIEGEGEGRPYEGTQTAKLKVTKGGPLPFAWDILSPQFMYGSKAYVKHPADIPDYLKLSFPEGFKWERVMNFEDGGVVTVTQDSSLQDGEFIYKVKLRGTNFPSDGPVMQKKTMGWEASSERMYPEDGALKGEIKQRLKLKDGGHYDAEVKTTYKAKKPVQLPGAYNVNIKLDITSHNEDYTIVEQYERAEGRHSTGGMDELYKGGGSGGGSDEVDGGGPAGEASSIPNREGKPIPNPLLGLGSTRTGEGVSTAVAQDFTIAAKHAIAVEANTGKILYEKDATQPVEIASITKLITVYLVYEALENGSITLSTPVDISDYPYQLTTNSEASNIPMEARNYTVEELLEATLVSSANSAAIALAEKIAGSEKDFVDMMRAKLLEWGIQDATVVNTTGLNNETLGDNIYPGSKKDEENKLSAYDVAIVARNLIKKYPQVLEITKKPSSTFAGMTITSTNYMLEGMPAYRGGFDGLKTGTTDKAGESFVGTTVEKGMRVITVVLNADHQDNNPYARFTATSSLMDYISSTFHHHHHH<br>PenTag/CCS/mCherry/PenTag/HisTag | Sequence coding for a truncated penicillin binding protein 3 coupled to mCherry via a linker containing a Casp3 cleavage site followed by a C-terminal second truncated penicillin binding protein 3 via a linker containing a Casp3 cleavage site with a C-terminal His-Tag (used as crosslinker in hydrogels) | This work |
| **pHM532** | MGVSTAVAQDFTIAAKHAIAVEANTGKILYEKDATQPVEIASITKLITVYLVYEALENGSITLSTPVDISDYPYQLTTNSEASNIPMEARNYTVEELLEATLVSSANSAAIALAEKIAGSEKDFVDMMRAKLLEWGIQDATVVNTTGLNNETLGDNIYPGSKKDEENKLSAYDVAIVARNLIKKYPQVLEITKKPSSTFAGMTITSTNYMLEGMPAYRGGFDGLKTGTTDKAGESFVGTTVEKGMRVITVVLNADHQDNNPYARFTATSSLMDYISSTFGGGSGGDEVDSAGSAGAYSKIRQPKLSDVIEQQLEFLILEGTLRPGEKLPPERELAKQFDVSRPSLREAIQRLEAKGLLLRRQGGGTFVQSSLWQSFSDPLVELLSDHPESQYDLLETRHALEGIAAYYAALRSTDEDKERIRELHHAIELAQQSGDLDA | Sequence coding for a truncated penicillin binding protein 3 coupled C-terminally to a Pyruvate dehydrogenase complex repressor via a linker containing a | This work |

| | | | |
|---|---|---|---|
| | ESNAVLQYQIAVTEAAHNVVLLHLLRCMEPMLAQNVRQNFEL<br>LYSRREMLPLVSSHRTRIFEAIMAGKPEEAREASHRHLAFIEEIL<br>LDRSREESRRERSLRRLEQRKNKGELNSKLEGKPIPNPLLGLDST<br>RTGGGGMASNDYTQQATQSYGAYPTQPGQGYSQQSSQPYGQQ<br>SYSGYSQSTDTSGYGQSSYSSYGQSQNTGYGTQSTPQGYGSTG<br>GYGSSQSSQSSYGQQSSYPGYGQQPAPSSTSGSYGSSSQSSSYG<br>QPQSGSYSQQPSYGGQQQSYGQQQSYNPPQGYGQQNQYNSSS<br>GGGGGGGGGGNYGQDQSSMSSGGGSGGGYGNQDQSGGGGSG<br>GYGQQDRGGGGGLNDIFEAQKIEWHEHHHHHH<br>PenTag/PdhR/IDR/aviTag/HisTag | caspase-3 cleavage site which is coupled C-terminal to an RNA-binding protein FUS with a C-terminal avitag and His-Tag (for solubility tag) | |
| **pHM535** | MHHHHHHGVSTAVAQDFTIAAKHAIAVEANTGKILYEKDATQ<br>PVEIASITKLITVYLVYEALENGSITLSTPVDISDYPYQLTTNSEA<br>SNIPMEARNYTVEELLEATLVSSANSAAIALAEKIAGSEKDFVD<br>MMRAKLLEWGIQDATVVNTTGLNNETLGDNIYPGSKKDEENK<br>LSAYDVAIVARNLIKKYPQVLEITKKPSSTFAGMTITSTNYMLE<br>GMPAYRGGFDGLKTGTTDKAGESFVGTTVEKGMRVITVVLNA<br>DHQDNNPYARFTATSSLMDYISSTFGGGSGGGSDEVDGGGPAG<br>EASSIPNREGKPIPNPLLGLGSTRTGEVSKGEEDNMAIIKEFMRF<br>KVHMEGSVNGHEFEIEGEGEGRPYEGTQTAKLKVTKGGPLPFA<br>WDILSPQFMYGSKAYVKHPADIPDYLKLSFPEGFKWERVMNFE<br>DGGVVTVTQDSSLQDGEFIYKVKLRGTNFPSDGPVMQKKTMG<br>WEASSERMYPEDGALKGEIKQRLKLKDGGHYDAEVKTTYKAK<br>KPVQLPGAYNVNIKLDITSHNEDYTIVEQYERAEGRHSTGGMD<br>ELYK<br>HisTag /PenTag/CCS/mCherry | Sequence coding for PenTag coupled N-terminally to mCherry via a linker containing a caspase cleavage site (for purification tag) | This work |
| **pCJL217** | MAYSKIRQPKLSDVIEQQLEFLILEGTLRPGEKLPPERELAKQFD<br>VSRPSLREAIQRLEAKGLLLRRQGGGTFVQSSLWQSFSDPLVEL<br>LSDHPESQYDLLETRHALEGIAAYYAALRSTDEDKERIRELHHA<br>IELAQQSGDLDAESNAVLQYQIAVTEAAHNVVLLHLLRCMEPM<br>LAQNVRQNFELLYSRREMLPLVSSHRTRIFEAIMAGKPEEAREA<br>SHRHLAFIEEILLDRSREESRRERSLRRLEQRKNKGELNSKLEGK<br>PIPNPLLGLDSTRTGGGGMASNDYTQQATQSYGAYPTQPGQGY<br>SQQSSQPYGQQSYSGYSQSTDTSGYGQSSYSSYGQSQNTGYGT<br>QSTPQGYGSTGGYGSSQSSQSSYGQQSSYPGYGQQPAPSSTSGS<br>YGSSSQSSSYGQPQSGSYSQQPSYGGQQQSYGQQQSYNPPQGY<br>GQQNQYNSSSGGGGGGGGGGNYGQDQSSMSSGGGSGGGYGN<br>QDQSGGGGSGGYGQQDRGGGGGLNDIFEAQKIEWHEHHHHH<br>H<br>PdhR/IDR/aviTag/ HisTag | Sequence coding for a Pyruvate dehydrogenase complex repressor (PdhR) coupled to N-terminal FUS and an avitag (for testing the solubility of this protein) | Unpublished |
| **pCVC022** | MHHHHHHGVSTAVAQDFTIAAKHAIAVEANTGKILYEKDATQ<br>PVEIASITKLITVYLVYEALENGSITLSTPVDISDYPYQLTTNSEA<br>SNIPMEARNYTVEELLEATLVSSANSAAIALAEKIAGSEKDFVD<br>MMRAKLLEWGIQDATVVNTTGLNNETLGDNIYPGSKKDEENK<br>LSAYDVAIVARNLIKKYPQVLEITKKPSSTFAGMTITSTNYMLE<br>GMPAYRGGFDGLKTGTTDKAGESFVGTTVEKGMRVITVVLNA<br>DHQDNNPYARFTATSSLMDYISSTFTLRKIVQQGDAYQDSKAP<br>VQDGKEDTVIAVAPEDIYLIERVGNQSSQSVQFTPDSKAIPAPLE<br>AGTVVGHLTYEDKDLIGQGYITTERPSFEMVADKKIEK<br>HisTag /PBP3 | | [1] |
| **pKJ57** | MMENTENSVDSKSIKNLEPKIIHGSESMDSGISLDNSYKMDYPE<br>MGLCIIINNKNFHKSTGMTSRSGTDVDAANLRETFRNLKYEVR<br>NKNDLTREEIVELMRDVSKEDHSKRSSFVCVLLSHGEEGIIFGT<br>NGPVDLKKITNFFRGDRCRSLTGKPKLFIIQACRGTELDCGIETD<br>SGVDDDMACHKIPVEADFLYAYSTAPGYYSWRNSKDGSWFIQ<br>SLCAMLKQYADKLEFMHILTRVNRKVATEFESFSFDATFHAKK<br>QIPCIVSMLTKELYFYHHHHHHH<br>Casp3/HisTag | | Unpublished, used in [2] |
| **pRK793** | MKTEEGKLVIWINGDKGYNGLAEVGKKFEKDTGIKVTVEHPD<br>KLEEKFPQVAATGDGPDIIFWAHDRFGGYAQSGLLAEITPDKAF | | [3] |

QDKLYPFTWDAVRYNGKLIAYPIAVEALSLIYNKDLLPNPPKT
WEEIPALDKELKAKGKSALMFNLQEPYFTWPLIAADGGYAFKY
ENGKYDIKDVGVDNAGAKAGLTFLVDLIKNKHMNADTDYSIA
EAAFNKGETAMTINGPWAWSNIDTSKVNYGVTVLPTFKGQPSK
PFVGVLSAGINAASPNKELAKEFLENYLLTDEGLEAVNKDKPL
GAVALKSYEEELAKDPRIAATMENAQKGEIMPNIPQMSAFWYA
VRTAVINAASGRQTVDEALKDAQTNSSSNNNNNNNNNNLGIEG
RGENLYFQGHHHHHHHGESLFKGPRDYNPISSTICHLTNESDGH
TTSLYGIGFGPFIITNKHLFRRNNGTLLVQSLHGVFKVKNTTTLQ
QHLIDGRDMIIIRMPKDFPPFPQKLKFREPQREERICLVTTNFQT
KSMSSMVSDTSCTFPSSDGIFWKHWIQTKDGQCGSPLVSTRDG
FIVGIHSASNFTNTNNYFTSVPKNFMELLTNQEAQQWVSGWRL
NADSVLWGGHKVFMVKPEEPFQPVKEATQLMNRRRRR
MalE/TEVS/HisTag/TEV

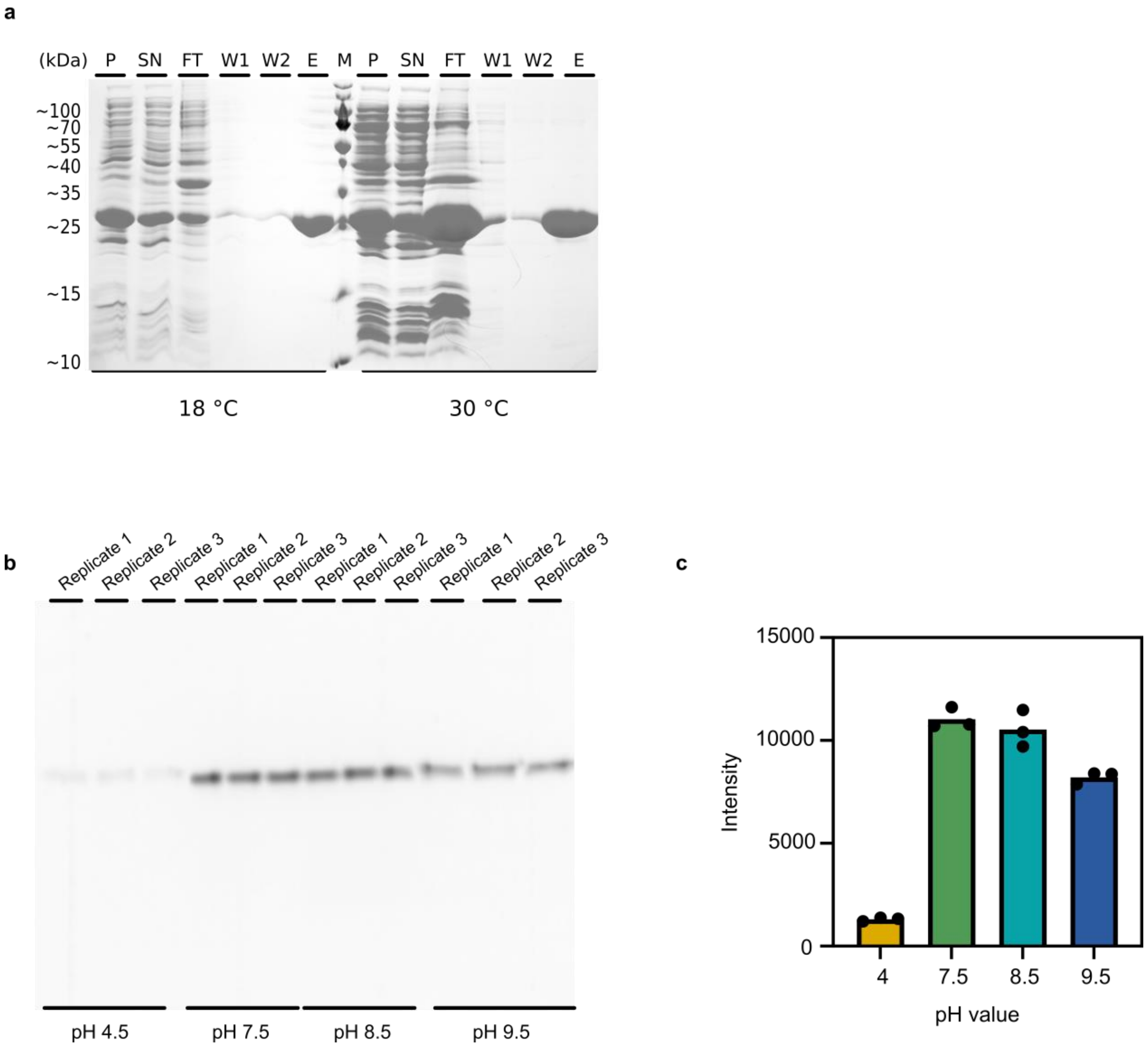


**Figure S1. SDS-PAGE analysis of PBP3-t purification by IMAC and analysis of PenTag-bocillin conjugate at different pH values. a,** Analysis of protein expression in *E. coli* and purification at two different temperatures by Ni-NTA. P, pellet; SN, supernatant; FT, flow-through; W1, wash fraction 1; W2, wash fraction 2; E, elution; M, marker. Expected size of the truncated protein is 30.2 kDa. **b**, Three replicates of bocillin-FL-conjugated PBP3-t at each pH condition were denatured and separated by SDS-PAGE. Both gels were run on a 15% (w/v) SDS-gel, 160 V. Visualization done by fluorescence filter. **c**, Intensity of the 3 replicates per sample analysed by ImageJ.

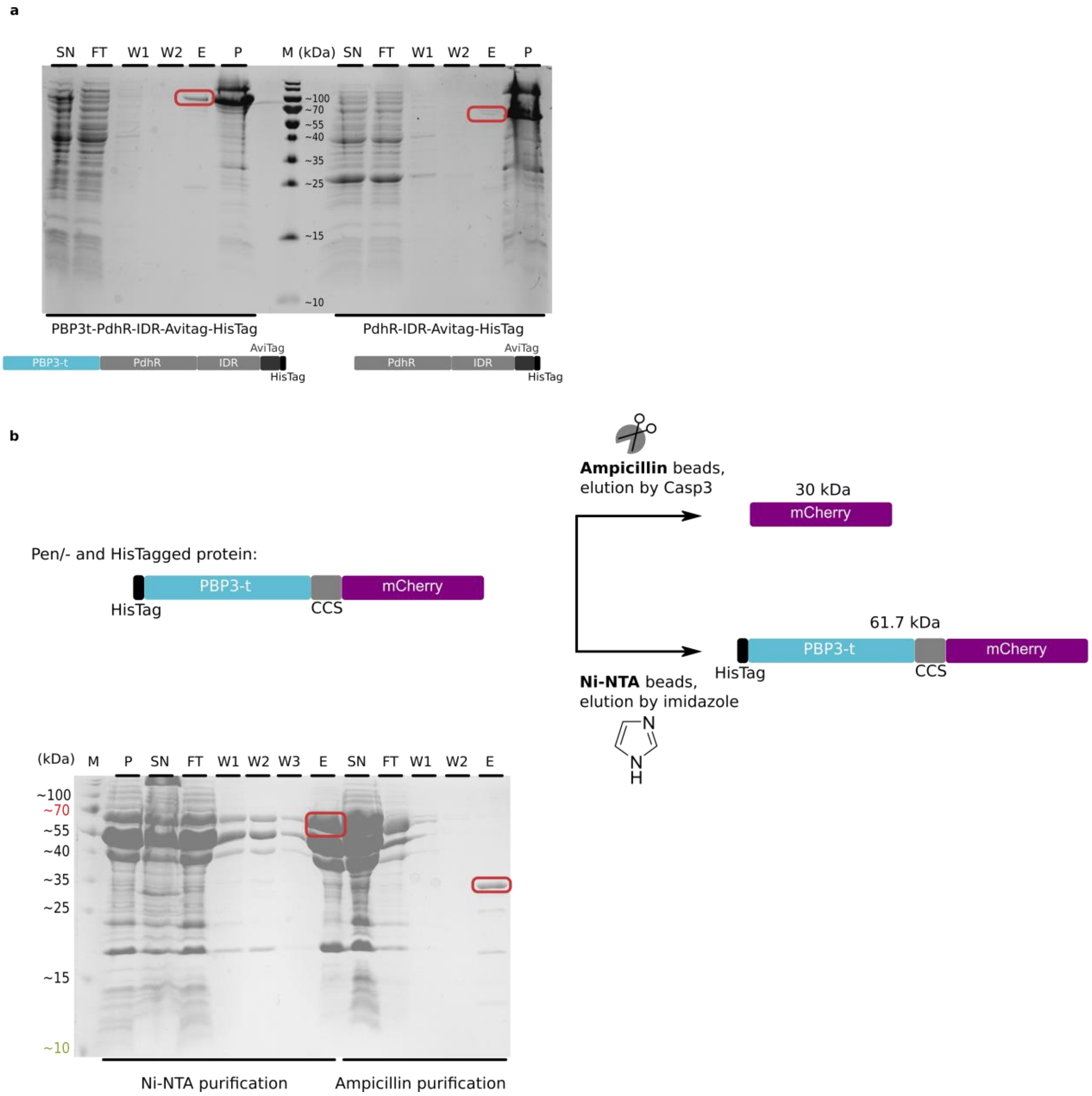


**Figure S2. PenTag technology as a solubility or purification tag. a,** Genetically fused PenTag as a solubility enhancing tag. SDS-PAGE analysis of protein expression in *E. coli* by Ni-NTA purification of PdhR-IDR with (left) and without (right) PBP3-t as solubility tag at the N-terminal. Expected sizes are 87.9 kDa and 56.7 kDa, respectively. A noticeable solubility enhancement can be observed in presence of fused PenTag. **b,** Purification analysis of HisTag-PBP3t-CCS-mCherry protein by Ni-NTA beads vs. ampicillin beads. Expected size of the cleaved protein from Ampicillin beads is 29.9 kDa. Whole protein purified by Ni-NTA beads is expected to be 61.7 kDa. P, pellet; SN, supernatant; FT, flow-through; W1, wash fraction 1; W2, wash fraction 2; E, elution; M, Marker. Both SDS-PAGE analyses performed on a 15% (w/v) SDS-gel.

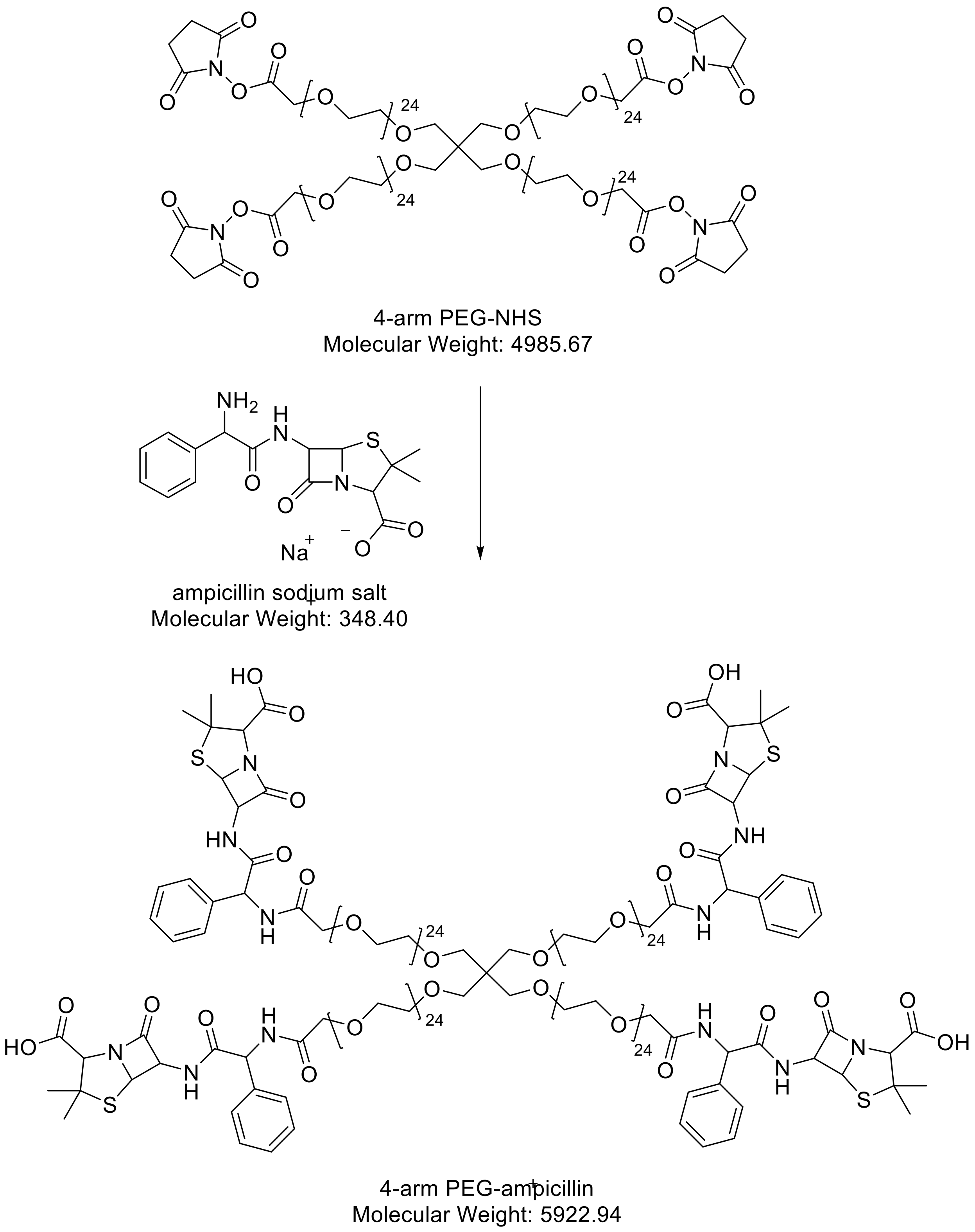


**Figure S3. Reaction scheme of ampicillin functionalization of 4-arm PEG-NHS.**

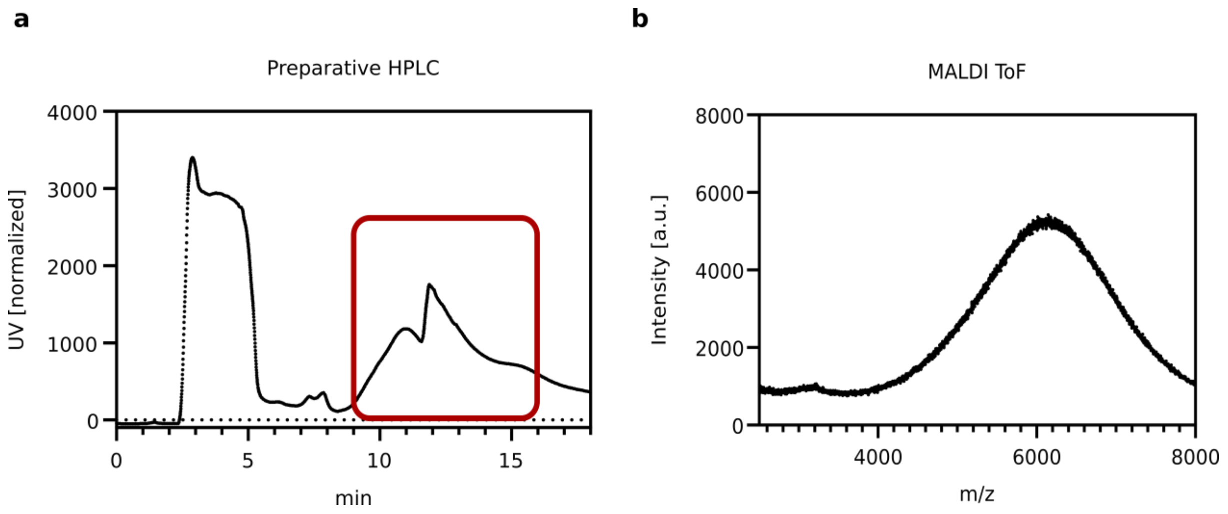


**Figure S4. Preparative reversed-phase HPLC diagram and MALDI-ToF results. a**, Preparative reversed-phase HPLC with collected fractions shown in red rectangle. Huge peak between 2-5 ml shows free excess ampicillin. **b**, MALDI-ToF shows a broad peak around 6000 g mol$^{-1}$ which is in agreement with the expected product. Broad peak is the result of the broad molecular weight distribution of reactant polymer and furthermore due to the fact that PEGs do not exist in aqueous solution as fully extended polymers and the polymer chains entangle to some degree.

a

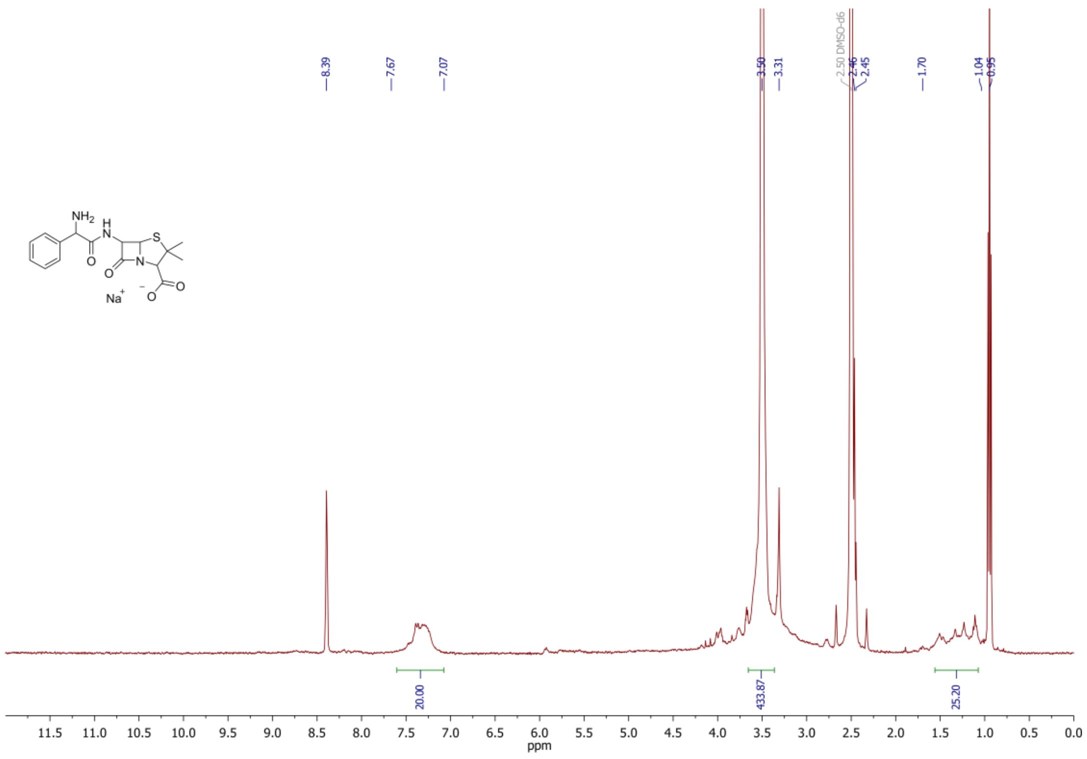

b

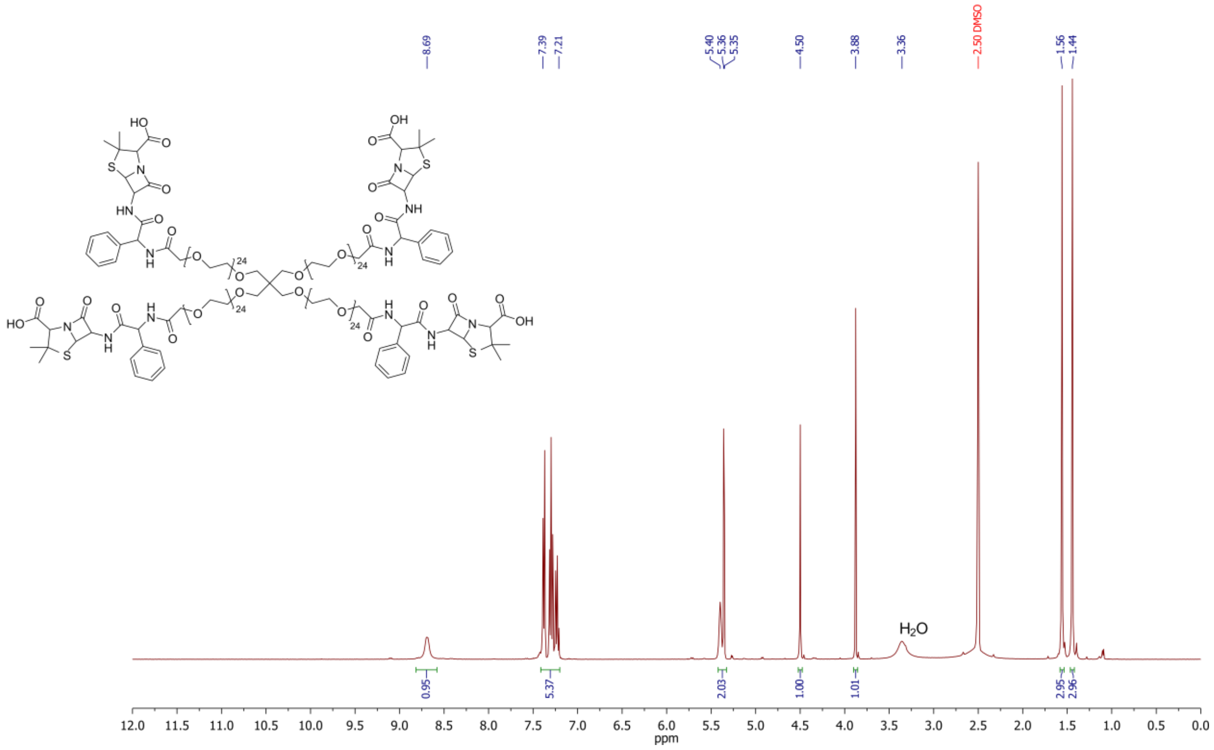

**Figure S5. NMR analysis of the reactant ampicillin, and reaction product. a**, Ampicillin: [1]H NMR (400 MHz, RT, DMSO-d$_6$): δ 8.69 (br. s, 1H, COOH acid), 7.39–7.21z (m, 5H, CH 1-benzene), 5.40 (m$_c$, 1H, CH propiolactam), 5.36 (dd, 1H, CH propiolactam), 4.50 (s, 1H, CH methine), 3.88 (s, 1H, CH methine), 3.36 (water), 1.56 (s, 3H, $CH_3$ methyl), 1.44 (s, 3H, $CH_3$ methyl). **b**, Functionalized polymer: [1]H NMR (400 MHz, RT, DMSO-d$_6$): δ 7.67–7.07 (m, 20H, CH 1-benzene), 3.62–3.34 (m, ~400H, $CH_2$

methylene-PEG), 1.58-1.08 (m, 24H, $CH_3$ methyl). In addition, residual water: 3.31 (s) and the counter-ion triethylammonium: 8.39 (s, 1H, N-$H^+$), 2.48 (q, 6H, $CH_2$), 0.95 (t, 9H, $CH_3$) were detected. Due to the nature of the non-defined PEG-chain-length, the insight by $^1$H-NMR is limited. The different chain lengths of the PEG-units result in different shifts for the end-group. Nevertheless, the average ratio of the characteristic groups (aromatic-, PEG $CH_2$- and $CH_3$-groups) proves the successful ampicillin functionalization.

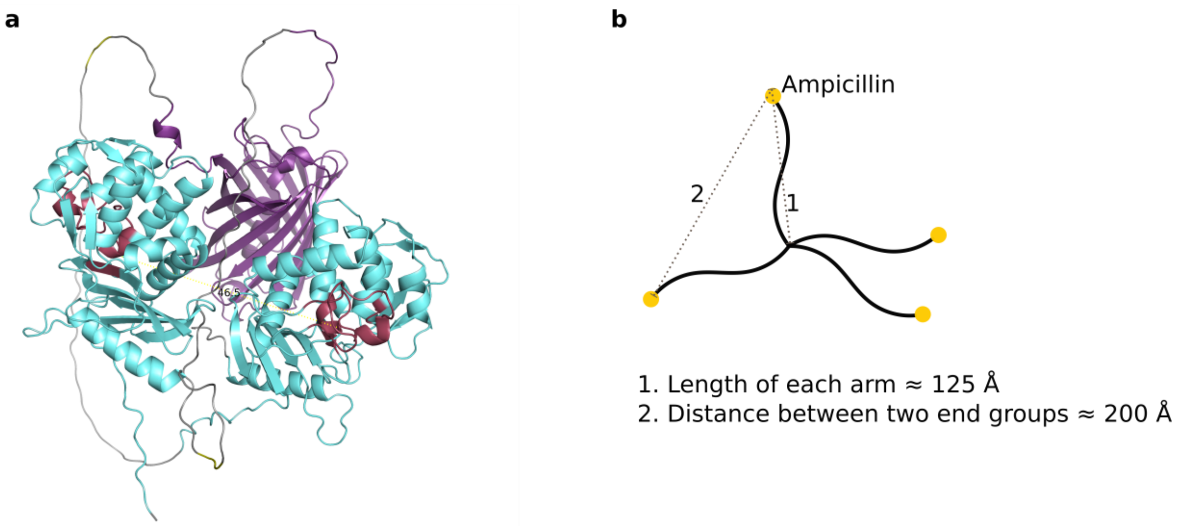


**Figure S6. Analysis of the distances between the crosslinking sites. a,** Protein structure shows both N-/ and C-terminal PBP3-t in light cyan and the mCherry protein in violet. Linkers are shown in grey with CCS in yellow. Active sites of the two PenTags shown in red, and the distance between them was measured by PyMOL corresponding to 46.5 Å. **b,** Approximate distances between the terminal ampicillin groups of each PEG-arm. 4-arm PEG with a molecular weight of 5000 kDa corresponds to 28 PEG repeating units per arm, each arm having a length of approximately 125 Å according to the manufacturer. With an assumed tetrahedral structure of the polymer, the distance between the end groups would be approximately 200 Å. However, entanglements of the polymer arms are a possibility for reduced distances between the end groups. Therefore, intramolecular bonds are a possibility.

**Table S2. Different ratios of PenTag : ampicillin-functionalized PEG-arm tested for hydrogels.** Only the last two conditions solidified as a hydrogel, whereas with the 1:8 ratio, mixing of the components were challenging due to direct gelation of the components on the pipette tips.

| PenTag : ampicillin-functionalized PEG-arm | 8:1 | 4:1 | 2:1 | 1:1 | 1:2 | 1:4 | 1:8 |
|---|---|---|---|---|---|---|---|
| Observation | Not solidified | Not solidified | Not solidified | Not solidified | Not solidified | Solidified | Solidified (difficult mixing) |

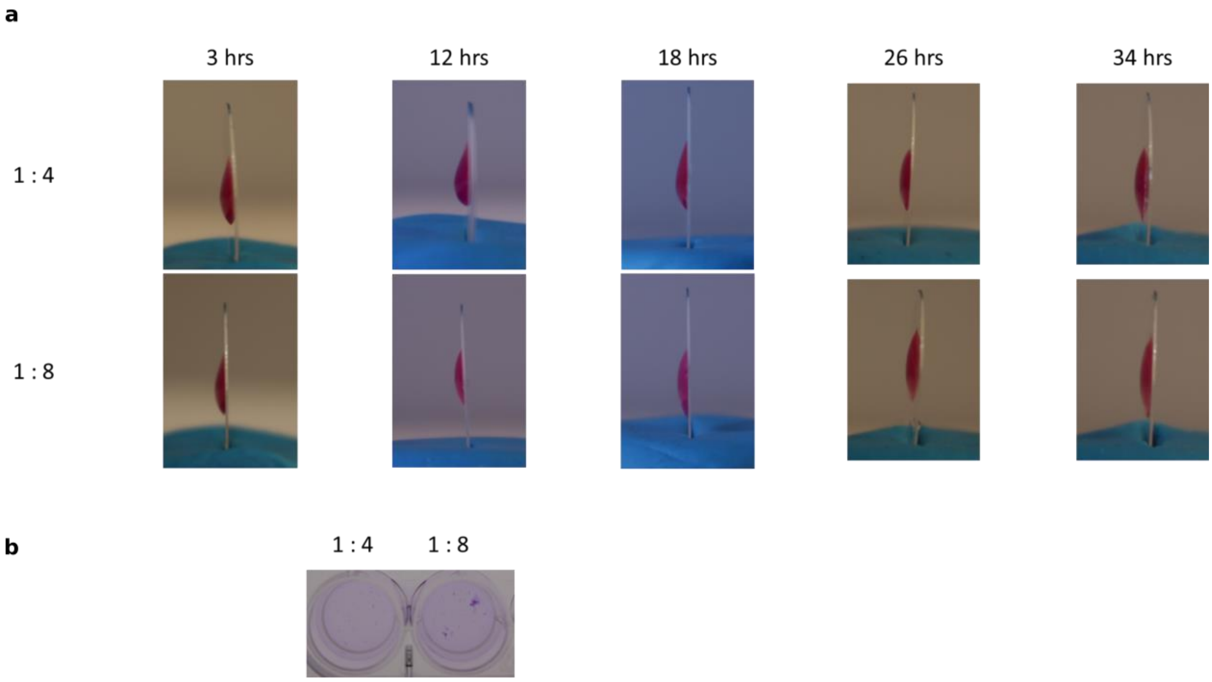


**Figure S7. Kinetics of gelation. a,** Hydrogels with PenTag : ampicillin-functionalized PEG-arm molar ratios of 1:4 and 1:8 were monitored over time. Hydrogels were prepared on separate (sigmacote-coated) round glass slides (1 cm diameter), as described in the experimental section. For observing the gelation process, each gel was vertically placed for studying the profile of the hydrogel at a specific time point. **b,** Hydrogels that looked symmetrical at 34 hours were transferred to PBS buffer and did not remain intact.

**a** **b**

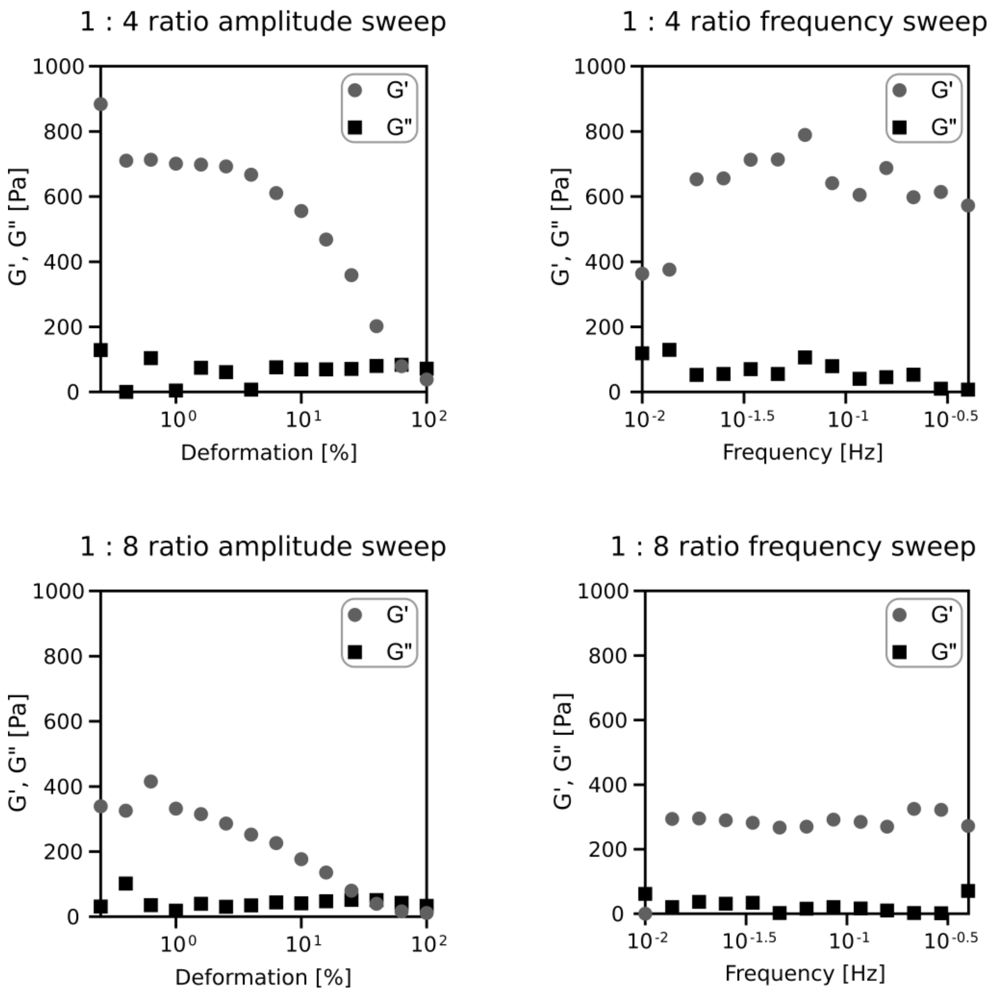


**Figure S8. Characterization of PenTag-hydrogels with PenTag : ampicillin-functionalized PEG-arm molar ratios of 1:4 and 1:8.** The storage (G') and loss (G") moduli were determined by small amplitude shear rheology via MCR301 rheometer (Anton Paar) with parallel plates at RT. The hydrogels were placed in between the plates (upper plate: 4 mm diameter, PP04, Anton Paar). **a,** a deformation range of 0.25 to 100% was measured at 0.3 Hz. **b,** at a constant deformation of 1%, a frequency range of 0.01 to 0.3 Hz was measured.

### Mathematical model description

#### 1. Mathematical model of the encoder

We developed a mathematical model to characterize, predict and optimize the behavior of a biomaterials-based encoder system. This model consists of ordinary differential equations (ODEs) describing the dynamic behavior of relevant system components. Enzymatic reactions were modeled via Michaelis Menten kinetics and partly reduced to mass action kinetic if applicable as described below.

The model scheme of the encoder system is shown in Figure S9a. mCherry is attached to beads by PBP3 and an anchor with two cleavage sites. These sites can be cleaved by TVMV (v1) or Casp3 (v2) resulting in the release of unbound mCherry. The TVMV cleavage site can be covered by the fluorescent protein and thus be party inaccessible for TVMV. These mCherry molecules can still be released by Casp3 (v3). Similar, GFP is attached to beads by a HaloTag and an anchor with two cleavage sites. Here, Casp3 (v4) as well as TEV (v5) can release GFP. Again, one cleavage site can be covered by fluorescent protein, this time it is the Casp3 cleavage site, providing additional GFP released only by TEV (v6). The model consists of 6 reactions with corresponding fluxes v1-v6 (Figure S9b). Given these fluxes, the resulting set of ODEs can be formulated (Figure S9c) to describe the dynamic behavior of the system.

a

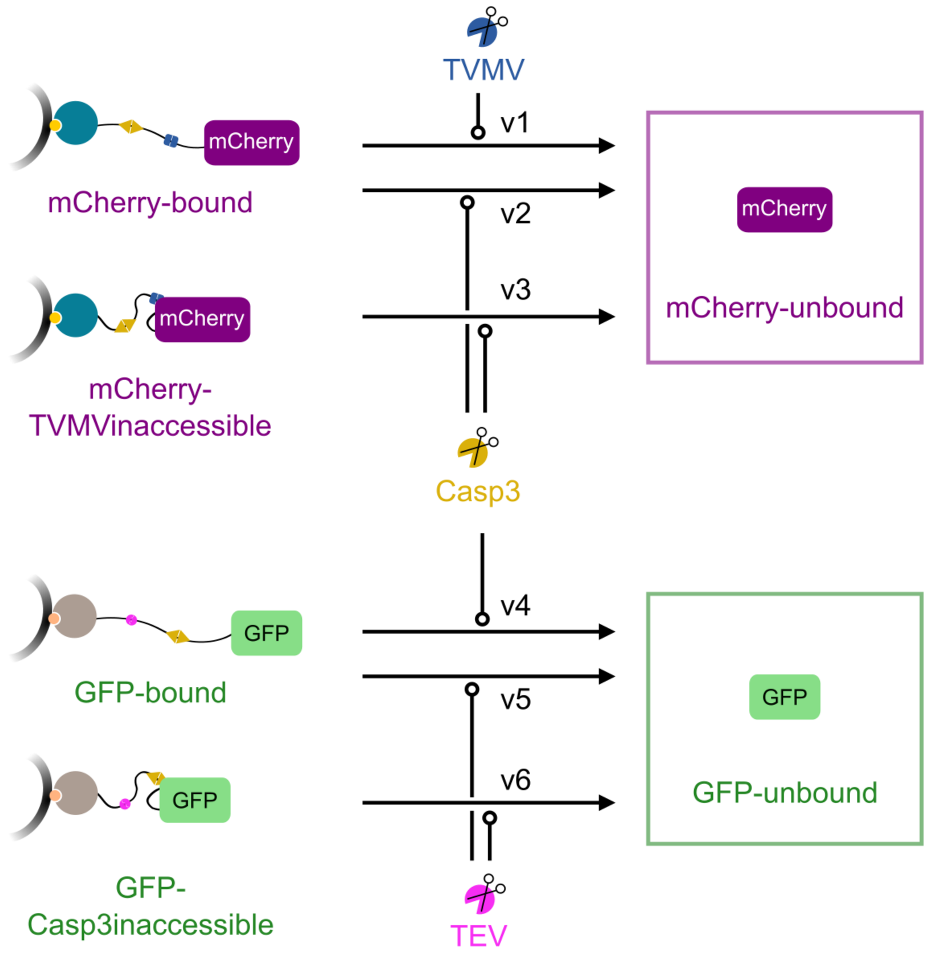


b

$$v_1 = \mathrm{TVMV} \cdot k\_\mathrm{releaseMC_{TVMV}} \cdot \frac{\mathrm{mCherry_{bound}}}{(\mathrm{Km_{TVMV}} + \mathrm{mCherry_{bound}})}$$

$$v_2 = \mathrm{Casp3} \cdot k\_\mathrm{releaseMC_{Casp3}} \cdot \mathrm{mCherry_{bound}}$$

$$v_3 = \mathrm{Casp3} \cdot k\_\mathrm{releaseMC_{Casp3}} \cdot \mathrm{mCherry_{TVMVinaccessible}}$$

$$v_4 = \mathrm{Casp3} \cdot k\_\mathrm{releaseGFP_{Casp3}} \cdot \frac{\mathrm{GFP_{bound}}}{(\mathrm{Km_{Casp3GFP}} + \mathrm{GFP_{bound}})}$$

$$v_5 = \mathrm{TEV} \cdot k\_\mathrm{releaseGFP_{TEV}} \cdot \frac{\mathrm{GFP_{bound}}}{(\mathrm{Km_{TEV}} + \mathrm{GFP_{bound}})}$$

$$v_6 = \mathrm{TEV} \cdot k\_\mathrm{releaseGFP_{TEV}} \cdot \frac{\mathrm{GFP_{Casp3inaccessible}}}{(\mathrm{Km_{TEV}} + \mathrm{GFP_{Casp3inaccessible}})}$$

c

$$\frac{d}{dt}\mathrm{mCherry_{bound}} = -v_1 - v_2$$

$$\frac{d}{dt}\mathrm{mCherry_{TVMVinaccessible}} = -v_3$$

$$\frac{d}{dt}\mathrm{mCherry_{unbound}} = v_1 + v_2 + v_3$$

$$\frac{d}{dt}\mathrm{GFP_{bound}} = -v_4 - v_5$$

$$\frac{d}{dt}\mathrm{GFP_{Casp3inaccessible}} = -v_6$$

$$\frac{d}{dt}\mathrm{GFP_{unbound}} = v_4 + v_5 + v_6$$

**Figure S9. Mathematical model of the Encoder. a,** Model scheme. mCherry is released by TVMV (v1) and Casp3 (v2, v3) respectively. GFP is released by Casp3 (v4) and TEV (v5, v6). **b,** Equations describing the reactions indicated in a. **c,** Overview of the ordinary differential equations (ODEs) of the encoder model.

### 2. Likelihood based parameter estimation and identifiability analysis

Model parameters were a priori unknown and were therefore estimated from experimental data using maximum likelihood estimation. Internal model states were linked to the experimental data by an observation function, introducing scaling and offset parameters. Moreover, we assumed a constant Gaussian error as measurement error, whose variance was estimated simultaneously with the scaling and offset as well as the dynamic model parameters.

To determine parameter uncertainties and thus the identifiability of the system we calculated parameter confidence intervals using the profile likelihood method[4]. These parameter profiles were also used for model reduction as previously described[5]. The reduction affected mainly the release of GFP by Casp3. Here, Michaelis Menten kinetics were reduced to mass action kinetics to render the system identifiable.

### 3. Implementation of single experiments

Four different experiments, characterizing the time-resolved as well as the dose-resolved release of GFP and mCherry, were used to calibrate the encoder model.

Experiment 1 and 2: Characterization of the mCherry OR gate

In these experiments, the mCherry OR gate, as outlined in the upper part of Figure S6a, was implemented to quantify the release of mCherry by TVMV and Casp3. First, the mCherry release was measured in dependence of different concentrations of TVMV and TEV. Second, the release was monitored for 85 minutes after stimulation with a fixed concentration of the two enzymes. Initial concentrations of TVMV and Casp3 were fixed to the respective applied concentrations and initials of mCherry-bound as well as the TVMV-inaccessible form of mCherry-bound were estimated as model parameters.

Experiment 2 and 3: Characterization of the GFP OR gate

In these experiments, the GFP OR gate, as outlined in the lower part of Figure S6a, was implemented to quantify the release of GFP in dependence of TEV and Casp3. First, its release was measured in dependence of different concentrations of TVMV and TEV. Second, the release was monitored for 85 minutes after stimulation with a fixed concentration of the two enzymes. Initial concentrations of TEV and Casp3 were fixed to the respective applied concentrations and initials of GFP-bound as well as the Casp3-inaccessible form of GFP-bound were estimated as model parameters.

### 4. Fitting process and results

The two OR gates were combined to the complete encoder system. All model parameters were estimated simultaneously from the experimental data. In total, the model comprises 20 parameters, including seven dynamic parameters, four initial parameters, seven scaling parameters and two error parameters. Log-transformation of parameters was used to ensure positivity and numerical stability. Estimated parameter values corresponding to the global optimum along with profile likelihood-derived confidence intervals are depicted in Figure S10 as direct output of the analysis and listed in Table S3 on the linear scale.

Compilation, numerical integration, fitting and optimization of the model was performed with the R based free available software dMod[6]. Deterministic multi-start optimization was performed using the trust region optimizer[7]. Out of in total 50 fits all converged to the same lowest minimum indicating the global optimum. The resulting model curves and data are shown in Figure 4e-h. Shaded bands correspond to the estimated standard deviation of the Gaussian error model.

**Table S3. Parameters of the feedforward and the feedback system.** Values are estimated by the maximum likelihood method. $\sigma^-$ and $\sigma^+$ show the 95 % point wise confidence intervals calculated by the profile likelihood method.

| Parameter | $\Theta_{opt}$ | $\sigma^-$ | $\sigma^+$ |
|---|---|---|---|
| $GFP_{bound}$ | 0.079925 | 0.062195 | 0.097606 |
| $GFP_{Casp3inaccessible}$ | 0.010275 | 0.00642 | 0.014606 |
| $k\_releaseGFP_{Casp3}$ | 6.341807 | 3.864945 | 14.89564 |
| $k\_releaseGFP_{TEV}$ | 1.235833 | 0.758114 | 2.870928 |
| $k\_releaseMC_{Casp3}$ | 16.59262 | 14.81907 | 18.5176 |
| $k\_releaseMC_{TVMV}$ | 0.01405 | 0.009953 | 0.023849 |
| $Km_{Casp3GFP}$ | 0.048157 | 0.01489 | 0.182699 |
| $Km_{TEV}$ | 0.049828 | 0.016307 | 0.177958 |
| $Km_{TVMV}$ | 0.042601 | 0.016917 | 0.114781 |
| $mCherry_{bound}$ | 0.098976 | 0.084061 | 0.114251 |
| $mCherry_{TVMVinaccessible}$ | 0.041167 | 0.033178 | 0.049876 |
| $offset\_GFP_{DR}$ | 244.1715 | 109.4077 | 380.8458 |
| $offset\_mCherry_{DR}$ | 343.0115 | 227.3357 | 459.9953 |
| $offset\_mCherry_{TC}$ | 299.2642 | 188.4611 | 412.3121 |
| $scale\_GFP_{DR}$ | 1.678714 | 1.371553 | 2.157832 |
| $scale\_GFP_{TC}$ | 1.563758 | 1.280908 | 2.007977 |
| $scale\_mCherry_{DR}$ | 1.665402 | 1.450165 | 1.950359 |
| $scale\_mCherry_{TC}$ | 1.760945 | 1.523785 | 2.07249 |
| sigma_GFP | 163.128 | 143.0451 | 188.5753 |
| sigma_mCherry | 220.8207 | 194.0895 | 254.5065 |

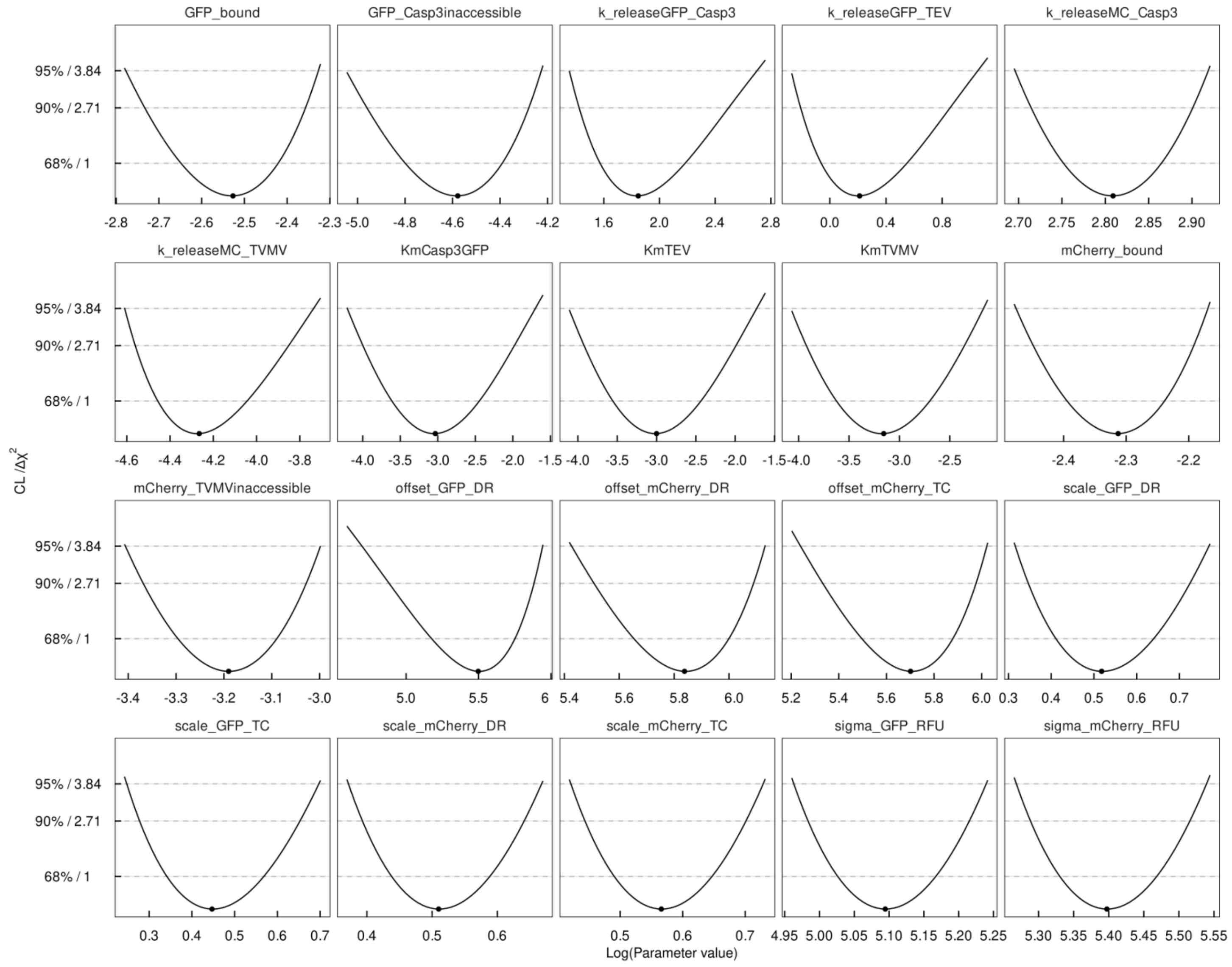


**Figure S10. Profile likelihood of estimated parameter values.** The profile likelihood is indicated by solid lines with the optimal parameter value marked by a dot. Confidence levels are indicated by dashed lines. Parameter values are depicted on the logarithmic scale.

### 5. Model predictions and validation

In order to optimize the encoder system to achieve a clear 4:2 encoder response after 30 minutes, optimal input concentrations of TVMV, TEV and Casp3 were determined by model simulations as 0.014 mg ml$^{-1}$ TEV, 0.8 mg ml$^{-1}$ TVMV, 0.006 mg ml$^{-1}$ Casp3 and 0.1 mg ml$^{-1}$ GFP and mCherry. Time-resolved predictions of the encoder system for these concentrations are shown in Figure S11a along with prediction confidence intervals. The TVMV/Casp3-inaccessible part of fluorescence protein was neglected and set to zero for this prediction as full accessibility was ensured in the setting of the additional experiment by significantly increasing the volume of the experiment.

To validate the model, dynamics of the full encoder system were also experimentally quantified after addition of predicted enzyme concentrations (Figure S11b). Dynamics of this additional one-pot experiment were in agreement with the model predictions and could thereby validate the model.

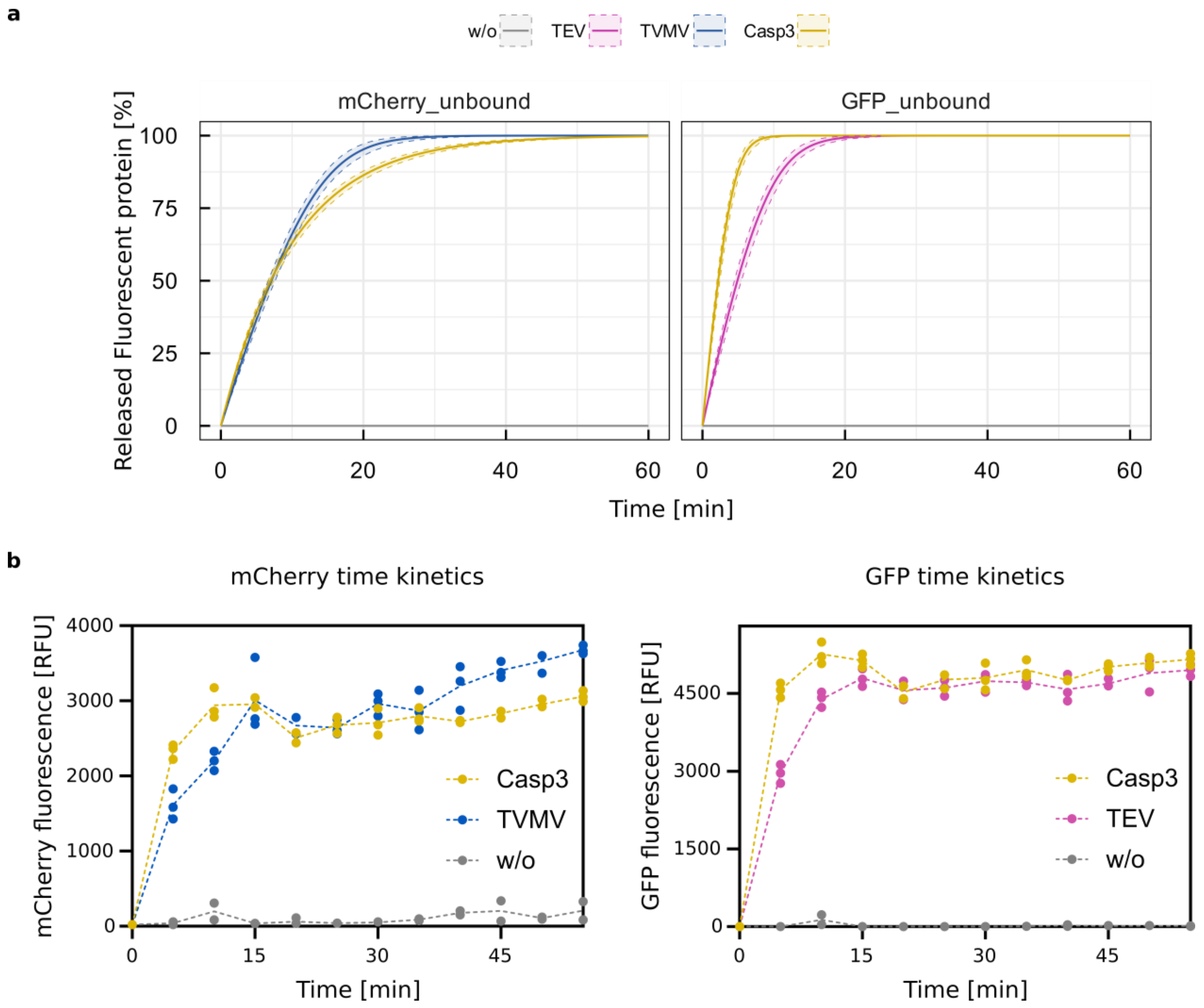


**Figure S 11. Prediction of the encoder dynamics and experimental time resolved measurements of the whole encoder one-pot set-up. a,** the release of mCherry and GFP was predicted for 0.014 mg ml$^{-1}$ TEV, 0.8 mg ml$^{-1}$ TVMV and 0.006 mg ml$^{-1}$ Casp3 respectively based on the calibrated encoder model. Model predictions are indicated as solid lines along with shaded error bands that describe the 1σ prediction confidence intervals. **b,** Raw data of the whole encoder system set-up. Samples measured in triplicates, and negative controls in duplicates. Dotted lines connect the mean of each condition.

**Table S4.** Overview of some of the covalent protein tagging systems in comparison to PenTag.[8–14]

| Protein Tag | Protein Size [kDa] | Coupling to | Advantages | Drawbacks |
|---|---|---|---|---|
| Halo-Tag | 33 | Haloalkane ligand | Rapid labelling ($10^6$ $M^{-1}$ $s^{-1}$)<br>High stability and specificity | High price<br>Synthetic haloalkane ligand |
| SNAP/CLIP-Tag | 19.4 | Alkylguanine/ benzylcytosine ligand | Small protein size<br>High stability and specificity | Lower labelling kinetics (2.8 x $10^4$ $M^{-1}$ $s^{-1}$)<br>High price<br>Synthetic complicated ligand<br>Purification necessary in the non-clickable SNAP-Tags |
| Catcher/Tag technology | ~12.3 | 13 aminoacid Tag | Parallel systems available (Spy, Snoop)<br>Diverse conditions (pH 5-10) and 4-37°C | High price<br>Peptide tag functionalization |
| Sortase A | ~25 | C-terminal LPXTG and N-terminal oligoglycine/alanine | High stability and specificity | High concentrations of oligoglycine needed<br>Challenging to reach complete conjugation<br>Removal of unreacted sortases |
| PenTag | 30.2 | Ampicillin<br>(can be further extended to other ß-lactam antibiotics) | Easy and cheap ligand<br>High stability and specificity<br>Spontaneous reaction | Abolished antibiotic activity of ampicillin during reaction |
